# Metabolic Coherence of the Mouse Brain

**DOI:** 10.64898/2026.05.07.723592

**Authors:** Zizhen Liu, Xin Ma, Roberto A. Ribas, Terrymar Medina, Sadi Quinones, Jihye Son, Lei Wu, Alison M. Ryan, Cameron Shedlock, Borhane Ziani, Vicenzo Barco-Caiaffa, Nidhi Rao, Kangjin Wong, Ann Titus, Reece Larson, Craig W. Vander Kooi, Navdeep S. Chandel, Matthew S. Gentry, Li Chen, Ramon C. Sun

## Abstract

The brain’s metabolic demands are well established, but how metabolism is coordinated across anatomically distinct regions remains poorly understood. Here, using matrix-assisted laser desorption/ionization (MALDI) imaging integrated with the Allen Brain Atlas and optimal transport-based computational analysis, we map the spatial metabolome across twelve major mouse brain divisions. We define an optimal-transport-derived inter-regional metabolite similarity metric and refer to it as metabolic coherence. This structure is largely preserved in an amyloid mouse model of Alzheimer’s disease despite widespread changes in individual metabolite and lipid levels. Individual metabolites and lipids shift in a coordinated manner across regions, sustaining inter-regional relationships even as absolute levels change in patterns indicative of mitochondrial dysfunction. To test whether the coherence metric is responsive to local intervention, we targeted the left hippocampus of mice from this model via lentiviral shHIF1α knockdown or neuronal AAV-mediated AOX expression. Both interventions were associated with metabolite normalization at the injection site. More importantly, normalization extended across distal regions sharing high metabolic similarity with the hippocampus and was accompanied by improved social memory in a single behavioral assay. Gene modulation and amyloid plaque reduction localized to the injection site.

## Introduction

The brain consumes roughly 20% of the body’s total energy despite comprising only 2% of its mass ^1^. Energy use is not uniform across brain regions, and aerobic glycolysis varies substantially across the cortex ^2^. Regional glycolytic patterns spatially correlate with amyloid-β deposition in individuals who later develop Alzheimer’s disease ^3^. Metabolic dysfunction is among the earliest and most consistent features of Alzheimer’s disease and related dementias, often preceding amyloid plaques and hyperphosphorylated tau by years to decades ^4,5,6,7^. Restoring brain glucose metabolism is sufficient to rescue cognition in mouse models of Alzheimer’s disease even when amyloid and tau pathology persist ^8^.

The brain operates as an interconnected network in which spatially distributed regions coordinate to support cognition and behavior.^1^ Neurodegenerative diseases follow this network architecture. Each major neurodegenerative syndrome shows selective vulnerability within a distinct intrinsic connectivity network, with Alzheimer’s disease preferentially targeting the default mode network.^9^ These intrinsic networks have been defined through functional connectivity analysis of resting-state fMRI, which reveals organized large-scale architectures including the default mode, frontoparietal, and salience networks.^10,11^ Whether metabolism is also organized at this network scale, and whether such organization carries functional significance for disease and intervention, has not been tested.

Inter-regional molecular similarity has been mapped most extensively through transcriptomics. The Allen Human Brain Atlas first showed that transcriptional profiles vary systematically by region and recapitulate known neuroanatomical boundaries.^12^ Richiardi and colleagues demonstrated that regions within the same resting-state functional network share correlated gene expression enriched for ion channels and synaptic genes, linking inter-regional transcriptomic similarity to synchronized neural activity.^13^ The BRAIN Initiative Cell Census Network has since produced whole-brain spatial transcriptomic atlases of the mouse, organizing thousands of cell types into a hierarchical taxonomy.^14,15^ Spatial transcriptomic mapping has identified molecular tissue regions across the mouse central nervous system that were previously unrecognized by classical neuroanatomy.^16^ MALDI mass spectrometry imaging (MALDI-MSI) extends this molecular mapping to metabolites, lipids, and glycans directly within tissue at near-cellular resolution.^17–21^ Trapped ion mobility and on-tissue derivatization workflows have revealed that brain lipid composition follows region-specific rather than global patterns.^22,23^ Unsupervised clustering of MALDI metabolite and lipid profiles recapitulates anatomical regions without prior spatial information, suggesting that metabolism itself tracks the organizational architecture of the brain.^24^ Whether brain metabolism is coordinated across anatomical regions as a network-level property of the brain, whether this coordination is preserved in Alzheimer’s disease, and whether it predicts distal metabolic response to a local intervention has not been directly tested.

Here, we define an optimal-transport-derived inter-regional metabolite similarity metric, which provides a network-level readout of brain metabolic organization, and refer to it as metabolic coherence. We then test whether a local hippocampal intervention in 5xFAD mice ^25,26^, using either lentiviral shHIF1α knockdown or neuronal AAV-mediated AOX expression, is associated with metabolite changes in distal regions of high metabolic similarity, and with improved social memory.

## Results

### MALDI mass spectrometry imaging reveals metabolic coherence across mouse brain regions

We previously developed the MetaVision3D pipeline for three-dimensional reconstruction of brain metabolome atlases from serial MALDI mass spectrometry imaging (MALDI-MSI) sections^27^ and the Sami platform for simultaneous profiling of metabolites, lipids, and glycans from single tissue sections.^24^ During these studies, we observed that individual metabolites and lipids consistently display consistent co-localization patterns across multiple anatomically distinct brain regions. This coordination is reminiscent of the transcriptomic co-expression architectures reported across the mammalian brain.^16–20^ The finding prompted us to systematically investigate whether brain metabolism exhibits a structured, inter-regional coordination that extends beyond individual molecular species.

MALDI-MSI of sagittal mouse brain sections revealed that individual molecular species exhibit distinct but spatially structured distributions across the brain. Lysophosphatidylethanolamine LPE(22:6) showed high abundance in the hippocampus and cerebral cortex with lower signal in the cerebellum, whereas lysophosphatidylethanolamine LPE(18:1) showed a complementary distribution with stronger signal in deeper structures and lower signal in the hippocampus and cerebellum (Figure 1A). Additional ion images of region-specific lipid species and a metabolite expression specificity heatmap are shown in Supplementary Figure 1. To extract region-level metabolite profiles from these imaging data, we developed a dual registration pipeline that aligns each MALDI section to the Allen Brain Atlas through both automated multi-stage transformation (rigid, affine, and B-spline) and manual refinement using QuickNII and VisuAlign (Figure 1B). Atlas-guided annotation-based segmentation produced anatomically faithful regional boundaries that intensity-based thresholding could not resolve, particularly in regions where metabolite gradients cross anatomical borders (Figure 1C). This registration framework enabled systematic quantification of metabolite distributions within each of the 12 major brain divisions and approximately 260 finer sub-regions defined by the Allen Brain Atlas, providing the anatomical foundation for all subsequent analyses. Three-dimensional renderings of the 12 major divisions and representative sub-regions are shown in Supplementary Figure 2.

**Figure 1.**
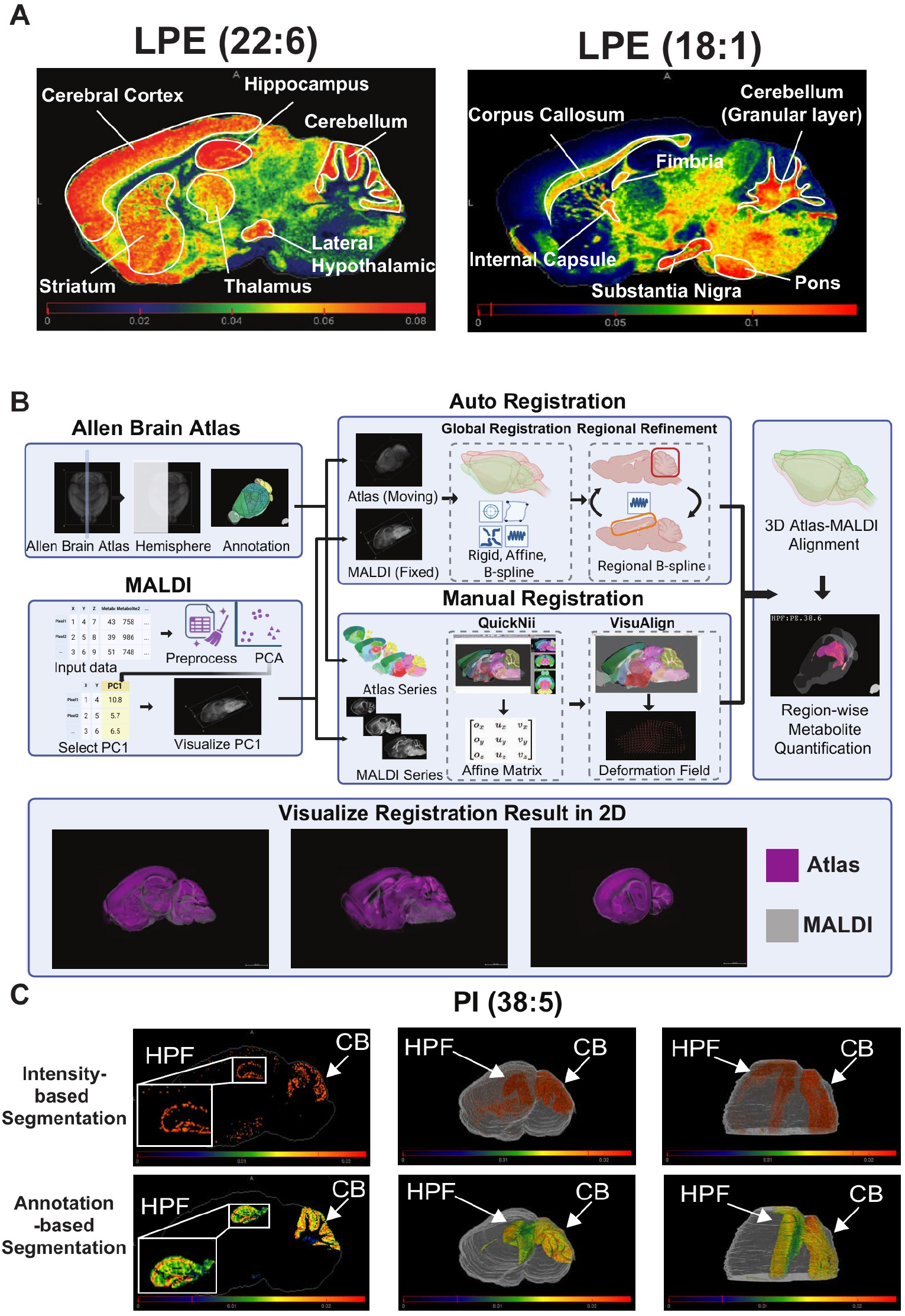
Spatial MALDI metabolomics framework for whole-brain metabolite mapping and atlas-guided region annotation. **(A)** Representative MALDI ion images illustrating region-specific spatial distribution of two lysophosphatidylethanolamine (LPE) species, LPE (22:6) and LPE (18:1), across sagittal mouse brain sections. Selected anatomical regions are labeled: Cerebral Cortex (Isocortex), Hippocampus (HPF), Cerebellum (CB), Striatum (STR), Thalamus (TH), Lateral Hypothalamic Area, Corpus Callosum, Fimbria, Granular Layer of Cerebellum, Internal Capsule, Substantia Nigra, and Pons (P). **(B)** Schematic overview of the MALDI-to-Allen Brain Atlas (CCFv3) registration and region-wise metabolite quantification pipeline. The Allen Brain Atlas including hemisphere volume and anatomical annotations, serves as the moving image for registration. MALDI imaging data undergo preprocessing and the first principal component (PC1) of the preprocessed metabolite matrix is used as the fixed image to guide registration, capturing dominant spatial variation in metabolite intensities while preserving original MALDI measurements. Up: Auto registration workflow. Sequential rigid, affine, and B-spline deformable transformations are applied using SimpleITK, followed by local refinement at anatomically complex regions using regional B-spline registration and Gaussian-smoothed displacement field integration. Down: Manual registration workflow. Global coarse alignment is performed in QuickNII by matching anatomical landmarks; local non-linear refinement is performed in VisuAlign using landmark-guided B-spline warping. The final registered atlas is used to assign each MALDI pixel to an anatomical region, enabling region-wise metabolite quantification at both coarse and fine region levels. **(C)** Comparison of intensity-based (top row) and annotation-based (bottom row) segmentation strategies applied to the PI (38:5). Annotation-based segmentation leverages registered atlas labels to define regions (e.g., HPF and CB), providing anatomically interpretable boundaries that are independent of individual metabolite signal intensity.

Using this framework, we extracted metabolite intensity distributions for each of the 12 major anatomical divisions from the three-dimensional sagittal MALDI-MSI dataset generated by MetaVision3D. To quantify inter-regional metabolic similarity, we computed pairwise inter-regional distances between weighted metabolite distributions using entropy-regularized optimal transport (Wasserstein)^28,29^ and converted these distances to similarity scores using a Gaussian kernel transformation (see Methods), generating a region-by-region metabolic coherence matrix (Figure 2A). This analysis revealed that metabolites maintain coherent co-localization patterns across brain regions with hierarchical organization visible in the similarity structure. Closely related regions such as the hippocampal formation (HPF) and isocortex exhibited high metabolic similarity, while phylogenetically and functionally distinct regions such as the cerebellum (CB) displayed more divergent metabolic profiles (Figure 2A–C). Three-dimensional visualization of the strongest inter-regional connections confirmed that metabolic similarity follows known neuroanatomical axes, with the HPF showing high average similarity to cortical, thalamic, and hypothalamic regions in this metric (Figure 2D and 2E). In contrast, the cerebellum formed a more isolated module with comparatively weaker connections to forebrain structures (Figure 2F and 2G).

**Figure 2.**
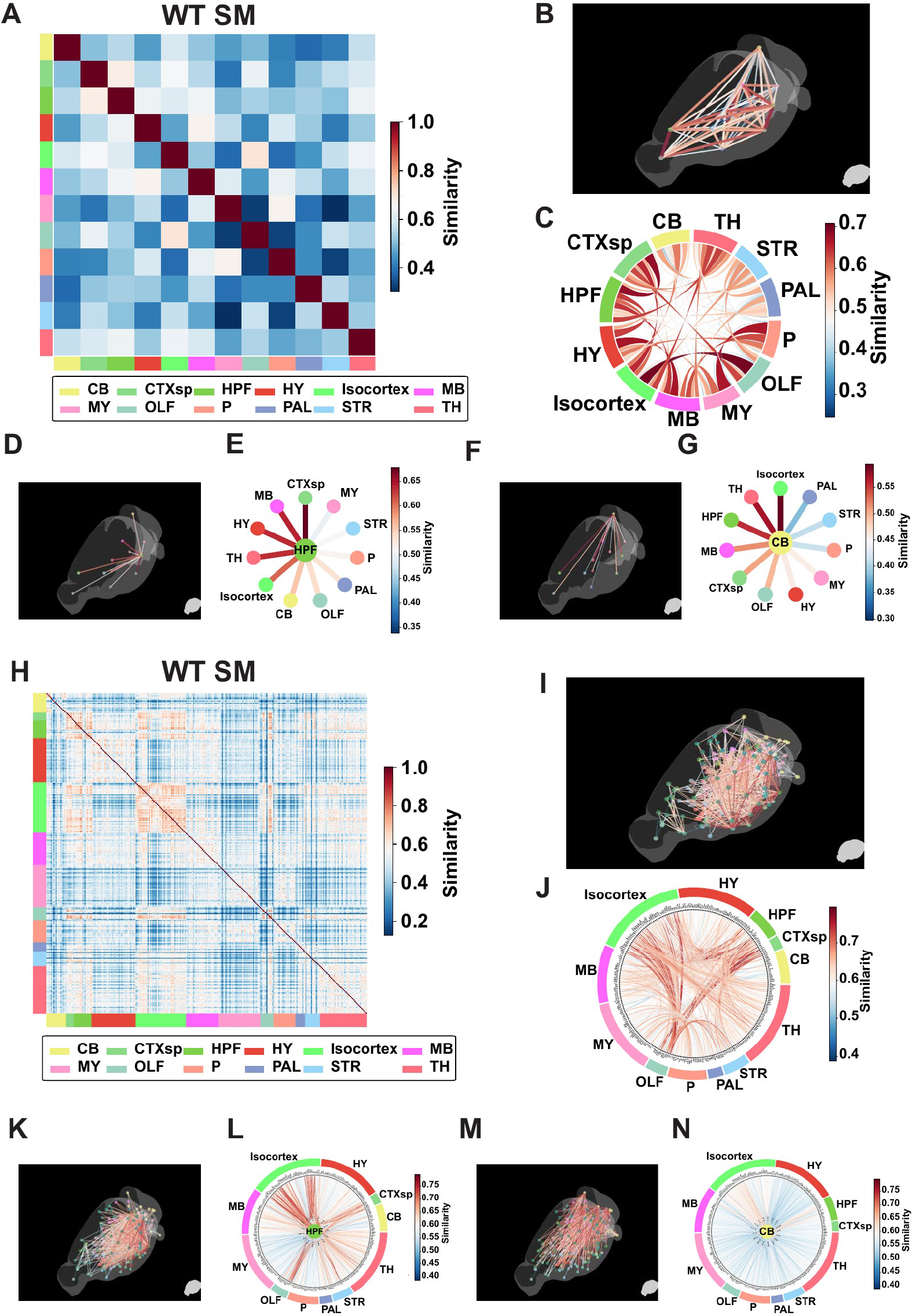
Whole-brain OT-based inter-(sub)region small molecule metabolomic coherence in wild-type mouse brain. **(A)** OT-based inter-region small molecule metabolomic coherence heatmap for a representative wild-type (WT) MALDI sample. Each entry represents the pairwise optimal transport (OT)-based similarity between the metabolite distributions of two anatomical regions (12 major Allen Brain Atlas structure sets). Similarity values range from 0 to 1. Regions: CB, cerebellum; CTXsp, cortical subplate; TH, thalamus; STR, striatum; OLF, olfactory areas; HPF, hippocampal formation; P, pons; HY, hypothalamus; PAL, pallidum; Isocortex; MB, midbrain; MY, medulla. **(B)** 3D spatial projection map of OT-based inter-region small molecule metabolomic coherence across major brain regions in WT mice. Lines connect region pairs; line color indicates OT-based similarity values. **(C)** Chord diagram summarizing OT-based inter-region small molecule metabolomic coherence across major brain regions in WT mice. Chord color and width reflect similarity values. **(D--E)** 3D spatial projection map (D) and radial similarity profile (E) centered on the hippocampal formation (HPF), illustrating OT-based inter-region small molecule metabolomic coherence relationships with all other major brain regions in WT mice. **(F--G)** 3D spatial projection map (F) and radial similarity profile (G) centered on the cerebellum (CB), illustrating OT-based inter-region small molecule metabolomic coherence relationships with all other major brain regions in WT mice. **(H)** OT-based inter-subregion small molecule metabolomic coherence heatmap for a representative WT sample. Subregions are grouped and color-coded by major structure set. **(I)** 3D spatial projection map of OT-based inter-subregion small molecule metabolomic coherence in WT mice. Lines connect subregion pairs; line color indicates similarity values. **(J)** Chord diagram summarizing OT-based inter-subregion small molecule metabolomic coherence in WT mice. Chord color and width reflect similarity values. **(K--L)** 3D spatial projection map (K) and chord diagram (L) at the subregion level centered on the HPF, illustrating OT-based inter-subregion small molecule metabolomic coherence relationships between HPF and all other brain regions in WT mice. **(M--N)** 3D spatial projection map (M) and chord diagram (N) at the subregion level centered on the CB, illustrating OT-based inter-subregion small molecule metabolomic coherence relationships between CB and all other brain regions in WT mice. In subregion-level visualizations (I--N), only the top 3 most similar subregion connections per subregion are displayed for visual clarity.

To assess whether this organization extends to finer anatomical scales, we repeated the analysis across approximately 260 sub-regions defined by the Allen Brain Atlas. The sub-regional similarity matrix recapitulated the parent-level structure while revealing additional within-region heterogeneity, with intra-regional sub-region pairs showing the highest similarity scores along the diagonal and graded inter-regional similarity reflecting known connectional architecture (Figure 2H–J). At this finer resolution, hippocampal formation sub-regions showed high average similarity to sub-regions across the isocortex, thalamus, and hypothalamus (Figure 2K and 2L). Cerebellar sub-regions again formed a relatively self-contained module (Figure 2M and 2N). The lipidome reproduced this coherence architecture at both region and sub-region scales (Supplementary Figure 3). We term this phenomenon, the coordinated maintenance of metabolic states across anatomically defined brain regions, metabolic coherence. Brain metabolism, in this framework, is organized as an inter-regional network. The observation that this coherence is maintained at both gross and fine anatomical resolution, and that the hippocampal formation shows the highest average similarity to other regions in this metric, raised an immediate question. Is metabolic coherence preserved under conditions of neurological disease, or does pathology disrupt the inter-regional coordination of brain metabolism?

### Metabolic coherence is preserved in an amyloid mouse model of Alzheimer’s disease despite widespread metabolite alterations

Having defined the inter-regional metabolic similarity structure in the wild-type mouse brain, we next asked whether this inter-regional coordination is disrupted under conditions of neurological disease. We examined the 5xFAD transgenic mouse model of Alzheimer’s disease, which harbors five familial AD mutations and develops progressive amyloid pathology, neuroinflammation, and behavioral deficits.^25,26^ Using the three-dimensional sagittal MALDI-MSI datasets generated by MetaVision3D (n = 1 per genotype),^27^ we computed optimal transport-based similarity matrices and found that the overall metabolic coherence structure was preserved in the 5xFAD brain. At the region level, pairwise similarity values between wild-type and 5xFAD were highly correlated (Spearman ρ = 0.859, p < 0.001), with permutation testing confirming that the observed correlation exceeded all values in the null distribution (Perm_p = 0.001) (Figure 3A–C). The distributions of region-level similarity values showed a moderate alteration between genotypes (Kolmogorov–Smirnov D = 0.227, p = 0.07) (Figure 3D and 3E), demonstrating that both the pattern and magnitude of region-level coherence were maintained in the 5xFAD brain.

**Figure 3.**
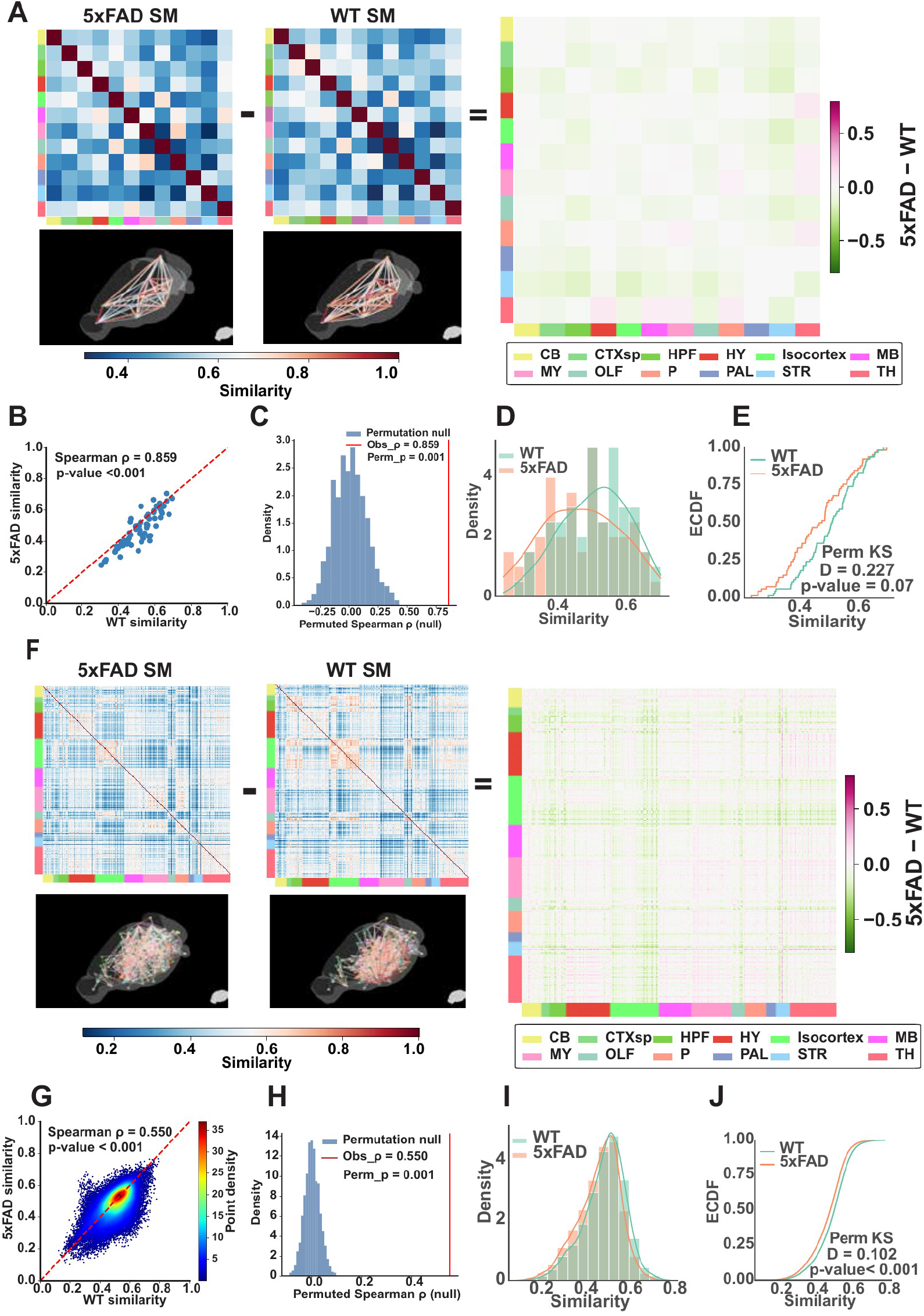
The global pattern of small molecule metabolomic coherence is highly conserved between 5xFAD and wild-type brains, with subtle alterations in regional similarity strength. **(A)** Side-by-side comparison of OT-based inter-region small molecule metabolomic coherence matrices for 5xFAD (left) and WT (right) at the major region level, and their difference matrix (5xFAD − WT). Below each matrix, the corresponding 3D spatial projection map is shown. **(B)** Scatter plot of paired OT-based inter-region small molecule similarity values from WT (x axis) versus 5xFAD (y axis) at the major region level. Each point represents one region pair. Spearman ρ = 0.859, p < 0.001. Dashed diagonal line indicates identity. **(C)** Null distribution of Spearman ρ under 1,000 node-label permutations at the major region level. Obs_ρ = 0.859, permutation p = 0.001, confirming that the OT-based inter-region small molecule similarity pattern is significantly preserved across genotypes. **(D)** Density distributions of OT-based inter-region small molecule similarity values from WT (green) and 5xFAD (orange) at the major region level. **(E)** ECDFs of OT-based inter-region small molecule similarity values from WT and 5xFAD at the major region level. Permutation-based Kolmogorov-Smirnov test: D = 0.227, p = 0.07, indicating a moderate alteration in the overall OT-based inter-region small molecule similarity distribution between 5xFAD and WT. **(F)** Side-by-side comparison of OT-based inter-subregion small molecule metabolomic coherence matrices for 5xFAD (left) and WT (right) at the subregion level, and their difference matrix (5xFAD − WT). Below each matrix, the corresponding 3D spatial projection map is shown. **(G)** Scatter plot of paired OT-based inter-subregion small molecule similarity values from WT (x axis) versus 5xFAD (y axis) at the subregion level. Each point represents one subregion pair; point color encodes density. Spearman ρ = 0.550, p < 0.001. Dashed diagonal line indicates identity. **(H)** Null distribution of Spearman ρ under 1,000 node-label permutations at the subregion level. Obs_ρ = 0.550, permutation p = 0.001, confirming that the OT-based inter-subregion small molecule similarity pattern is significantly preserved across genotypes. **(I)** Density distributions of OT-based inter-subregion small molecule similarity values from WT (green) and 5xFAD (orange) at the subregion level. **(J)** ECDFs of OT-based inter-subregion small molecule similarity values from WT and 5xFAD at the subregion level. Permutation-based Kolmogorov-Smirnov test: D = 0.102, p < 0.001, indicating a statistically significant yet modest alteration in the overall OT-based inter-subregion small molecule similarity distribution between 5xFAD and WT.

At the subregion level, the coherence architecture was similarly preserved. Pairwise similarity across approximately 260 sub-regions showed significant concordance between wild-type and 5xFAD (Spearman ρ = 0.550, p < 0.001), with permutation testing confirming significance (Perm_p = 0.001) (Figure 3F–H). Although the Kolmogorov–Smirnov test detected a statistically significant distributional shift (D = 0.102, p < 0.001) (Figure 3I and 3J), the effect size remained modest, reflecting the large number of pairwise comparisons at this resolution. The subregion-level analysis therefore reinforced the region-level finding. The network-level coordinated inter-regional organization of brain metabolism is maintained even in the presence of widespread metabolic perturbation. The full 5xFAD coherence panels at region and sub-region resolution are shown for the metabolome in Supplementary Figure 4 and for the lipidome in Supplementary Figure 5. The full statistical comparison of optimal transport-based coherence between 5xFAD and wild-type lipid samples at both region and subregion resolution is shown in Supplementary Figure 7, and a parallel analysis using a complementary Pearson correlation-based similarity measure is shown in Supplementary Figure 6.

Since the 3D sagittal MALDI dataset comprised n = 1 per genotype, we validated the preservation finding in n = 3 wild-type and n = 3 5xFAD coronal samples to establish biological reproducibility. We profiled the coronal sections by MALDI-MSI and registered them to the Allen Brain Atlas. Region-level OT similarity matrices for each replicate revealed consistent coherence patterns within and across genotypes (Figure 4A). The mean optimal transport similarity was not significantly different between wild-type and 5xFAD brains (Δ = 0.007, 95% CI [−0.021, 0.038], permutation p = 0.655) (Figure 4B), and permutation testing confirmed that within-genotype and between-genotype similarity structures were statistically indistinguishable (Obs_T = 0.020, Perm_p = 0.3) (Figure 4C). Pairwise Spearman correlation of group-mean similarity matrices demonstrated strong preservation of coherence architecture (ρ = 0.958, p < 0.001) (Figure 4D). Region-specific analysis revealed that the magnitude of similarity differences varied modestly across brain areas (Figure 4E). This independent validation in a different plane and cohort confirms that metabolic coherence is not an artifact of a single dataset or sectioning orientation. This pattern is consistent across anatomical planes and biological replicates.

**Figure 4.**
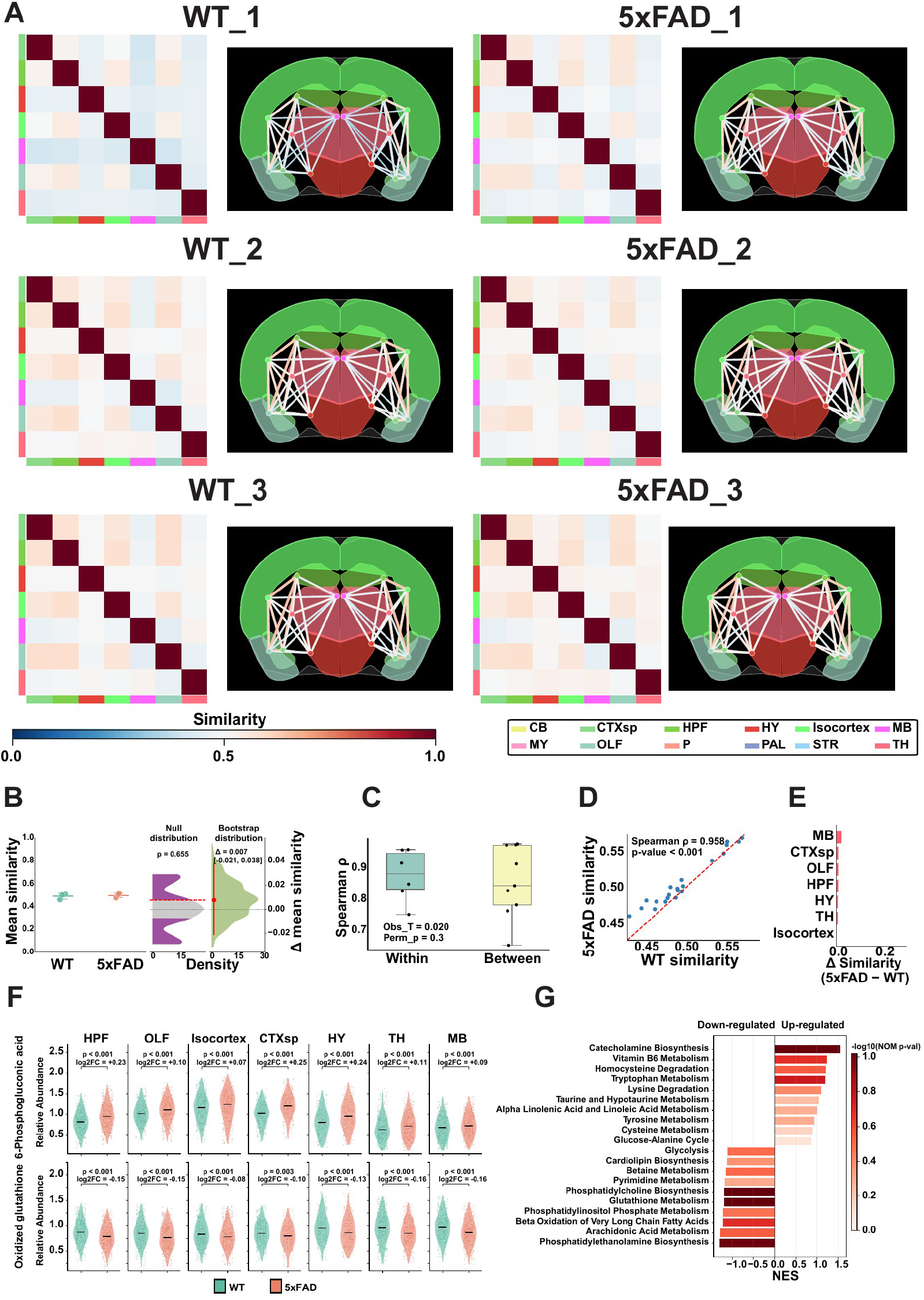
OT-based inter-region metabolomic coherence is largely preserved in 5xFAD and wild-type brains at 2D section level. **(A)** OT-based inter-region metabolomic coherence matrices and corresponding 2D spatial projection maps for three WT (left) and three 5xFAD (right) coronal brain sections at matched anatomical depths. Similarity values range from 0 to 1. **(B)** Gardner-Altman plot comparing mean OT-based inter-region metabolomic coherence between WT (n = 3) and 5xFAD (n = 3) samples. Left, individual data points showing sample-level mean similarity per group. Right, bootstrap distribution (10,000 replicates) of the mean difference (Δ = 0.007; 95% CI [−0.021, 0.038]; permutation p = 0.655), indicating no significant global shift in mean OT-based inter-region metabolomic coherence between genotypes. **(C)** Spearman rank correlation analysis of within-group versus between-group pairwise matrix concordance. Within-group Spearman ρ values (WT--WT and 5xFAD--5xFAD pairs) are compared to between-group ρ values (WT--5xFAD pairs). Obs_T = 0.020, permutation p = 0.3, indicating no significant difference in matrix concordance within versus between genotypes. **(D)** Scatter plot of paired OT-based inter-region similarity values from WT (x axis) versus 5xFAD (y axis). Each point represents one region pair. Spearman ρ = 0.958, p < 0.001. Dashed diagonal line indicates identity. **(E)** Region-level ΔSimilarity (5xFAD − WT) showing minimal per-region changes in mean OT-based inter-region metabolomic coherence across major brain regions. **(F)** Region-wise relative abundance of representative metabolites in 5xFAD (orange) versus WT (green) across individual brain regions. Top, 6-phosphogluconic acid, consistently upregulated in 5xFAD across all examined regions (log_2_FC range: +0.07 to +0.25; p < 0.001 for all regions). Bottom, oxidized glutathione, consistently downregulated in 5xFAD across all examined regions (log_2_FC range: −0.08 to −0.16; p ≤ 0.003 for all regions). **(G)** Metabolite set enrichment analysis (MSEA) comparing 5xFAD (n = 3) and WT (n = 3) sections. Metabolites were ranked by signal-to-noise statistic between groups and tested against SMPDB pathway gene sets using 1,000 phenotype permutations. Bar length, normalized enrichment score (NES); bar color, −log_10_(p value).

The preservation of coherence did not reflect an absence of metabolic change. Individual metabolite distributions showed significant alterations between genotypes, including increased 6-phosphogluconic acid and altered oxidized glutathione levels across multiple brain regions (Figure 4F). Metabolite set enrichment analysis revealed coordinated downregulation of catecholamine biosynthesis, vitamin B6 metabolism, and tryptophan metabolism alongside upregulation of phosphatidylethanolamine remodeling, pyrimidine metabolism, and glutathione metabolism in 5xFAD brains (Figure 4G). These data demonstrate that metabolic coherence is maintained not because the 5xFAD brain is metabolically unchanged, but rather because individual metabolites shift in a coordinated manner across regions, preserving inter-regional relationships even as absolute levels change. Per-sample metabolite Log2FC heatmaps for each WT–5xFAD replicate pair are shown in Supplementary Figure 13. The violin plots of individual metabolites across brain regions (Figure 4F) reveal that although the magnitude of each metabolite changes between genotypes, the relative pattern across regions is maintained: regions that are metabolically similar in wild-type remain similar in 5xFAD. This coordinated shifting of metabolite levels across regions is what we capture with the coherence metric. Analysis of metabolite spatial stability across anatomical axes (Supplementary Figures 8–11) and reproducibility of inter-regional coherence across independent coronal sections at matched depths (Supplementary Figure 12) confirmed that metabolic coherence was maintained through the three-dimensional extent of the brain. The inter-regional similarity structure is reproducible across anatomical planes and biological replicates in both wild-type and 5xFAD mice, even as individual metabolite levels change substantially.

### Targeted HIF1α knockdown in the hippocampus is associated with distal metabolite normalization and improved social memory

The preservation of metabolic coherence in 5xFAD brains despite widespread metabolite alterations raised an important question. If inter-regional metabolic states are coordinately maintained, could a local hippocampal intervention be associated with metabolite changes in distal regions sharing high metabolic similarity with the hippocampus? To identify candidate molecular targets, we performed whole-brain proteomics on wild-type and 5xFAD mice. Targeted analysis of metabolic enzymes revealed significant enrichment of oxidative phosphorylation, glycolysis, and tricarboxylic acid (TCA) cycle pathways in the 5xFAD brain (Figure 5A and 5B). This glycolytic shift is consistent with prior observations that brain regions with elevated aerobic glycolysis are preferentially vulnerable to amyloid pathology.^3^ Multiple glycolytic enzymes, identified as differentially expressed, are established transcriptional targets of hypoxia-inducible factor 1α (HIF1α), a master regulator of the metabolic switch from oxidative phosphorylation to glycolysis^30^ (Figure 5C and 5D). Immunofluorescence confirmed that HIF1α protein levels were significantly elevated in excitatory neuronal cell bodies of the CA1–CA3 subfields of the 5xFAD hippocampus compared to wild-type controls (p = 0.0463) (Figure 5E). These findings suggested that HIF1α-driven glycolytic reprogramming contributes to the metabolic alterations observed in the 5xFAD hippocampus and that reducing HIF1α activity could normalize hippocampal metabolism.

**Figure 5.**
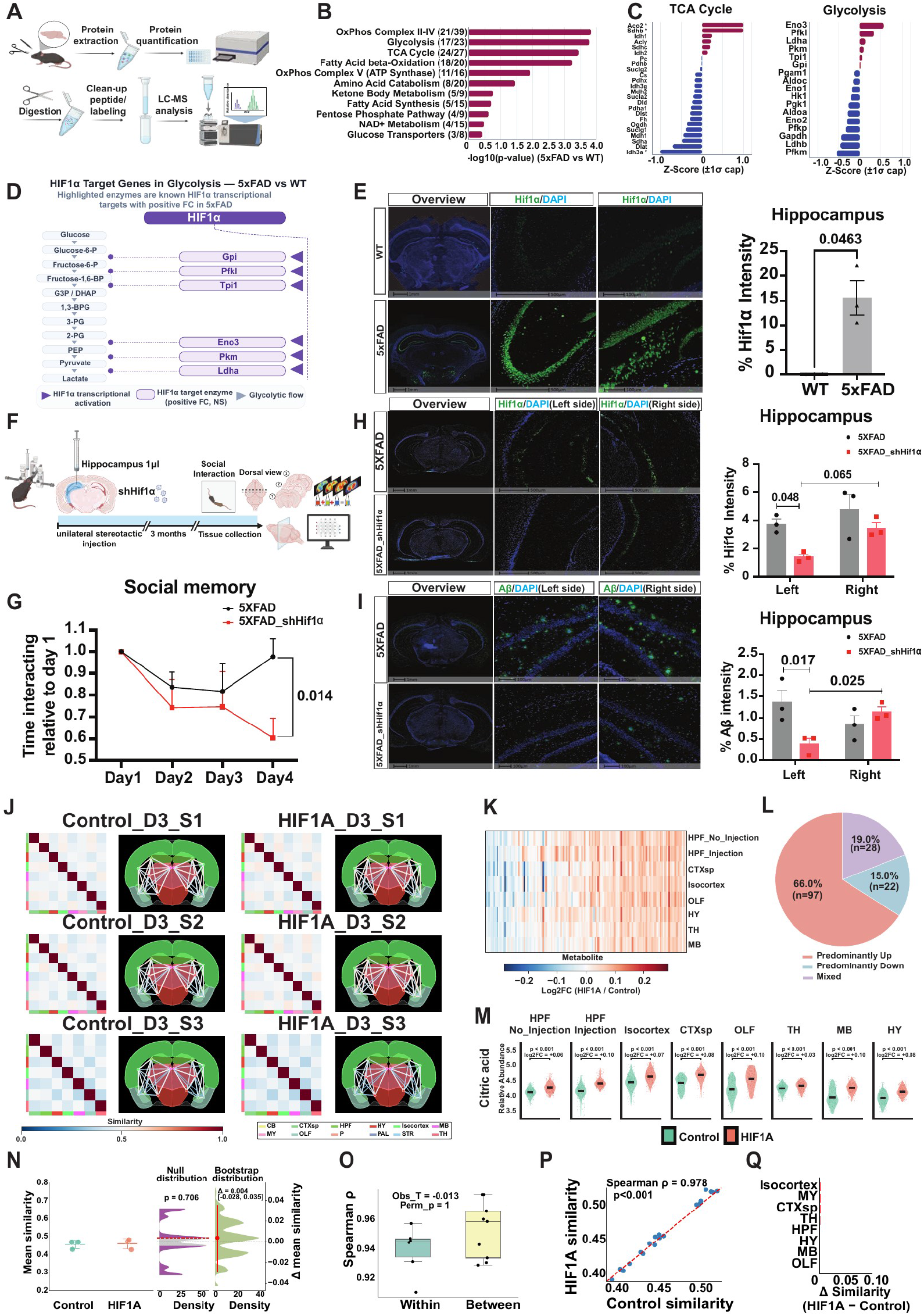
Proteomics Reveals HIF-1α-Driven Metabolic Reprogramming in 5xFAD Brain, and shHIF-1α Reduces Amyloid Pathology and Social Memory Deficits. **(A)** Schematic of proteomics workflow from 5xFAD and wild-type (WT) brain tissue. **(B)** Pathway enrichment analysis of differentially expressed membrane proteins in 5xFAD versus WT. Bars represent −log10(p-value) for each metabolic pathway. **(C)** Z-score-normalized protein abundance (±1σ cap) for TCA cycle (left) and glycolysis (right) enzymes in 5xFAD versus WT. **(D)** Schematic of hypoxia-inducible factor 1-alpha (HIF-1α) transcriptional targets within the glycolytic pathway. Enzymes with positive fold change (FC) in 5xFAD proteomics data are highlighted (purple boxes). **(E)** Representative immunofluorescence (IF) images of HIF-1α (green) and DAPI (blue) in coronal hippocampal sections from WT (top) and 5xFAD (bottom) mice. Scale bars, [500 μm (left) and 100 μm (right)]. Right: Quantification of HIF-1α fluorescence intensity in the hippocampus. Data are presented as mean ± SEM; n = 3 mice/group; [*p = 0*.*0463, two-tailed t-test*. **(F)** Schematic of unilateral hippocampal injection of shHif1α (1 μL) in 5xFAD mice. Social interaction testing was performed 3 months post-injection. **(G)** Social memory assessment. Interaction time relative to Day 1 is plotted across four consecutive days for 5xFAD-Lenti-shScrambled control (black) and 5xFAD-Lenti-shHIF1α (red) groups. Data are presented as mean ± SEM; n = 5-6 mice/group; [*p = 0*.*014, two-way ANOVA with multiple comparisons*. **(H)** Representative IF images of HIF-1α (green) and DAPI (blue) in the hippocampus of 5xFAD (top) and 5xFAD_shHif1α (bottom) mice. Scale bars, [500 μm]. Right: Quantification of HIF-1α fluorescence intensity in the hippocampus. Data are presented as mean ± SEM; n = 3 mice/group; [*p = 0*.*048 (left), p = 0*.*065 (right), two-tailed t-test*. **(I)** Representative IF images of amyloid-beta (Aβ; green) and DAPI (blue) in the hippocampus of 5xFAD (top) and 5xFAD_shHif1α (bottom) mice. Scale bars, [100 μm]. Right: Quantification of Aβ plaque intensity (% Aβ intensity) in the left and right hippocampus. Data are presented as mean ± SEM; n = 3 mice/group; [*p = 0*.*017 (left control V left shHif1α), p = 0*.*025 (shHif1 mice injection site V opposite side), two-tailed t-test*. **(J)** OT-based inter-region metabolomic coherence matrices and corresponding 2D spatial projection maps for 5xFAD Control (left) and 5xFAD HIF1A (right) coronal brain sections at anatomical depth 3. n = 3 sections per depth per group. **(K)** Heatmap of per-region metabolite log_2_FC (HIF1A/Control) across brain regions. Rows, brain regions; columns, metabolites. **(L)** Pie chart summarizing the directional consistency of metabolite abundance changes across regions. Metabolites are classified as Predominantly Up (>2/3 of regions increased; 66.0%, n = 97), Predominantly Down (>2/3 of regions decreased; 15.0%, n = 22), or Mixed (neither criterion met; 19.0%, n = 28). **(M)** Region-wise relative abundance of citric acid in HIF1A (orange) versus Control (green) across brain regions. Citric acid is consistently upregulated in HIF1A across all examined regions (log_2_FC range: +0.03 to +0.10; p < 0.001 for all regions). **(N)** Gardner-Altman plot comparing mean OT-based inter-region metabolomic coherence between Control (n = 3) and HIF1A (n = 3) samples. Left, individual data points showing sample-level mean similarity per group. Right, bootstrap distribution of the mean difference (Δ = 0.004; 95% CI [−0.028, 0.035]; permutation p = 0.706), indicating no significant global shift in mean OT-based inter-region metabolomic coherence between conditions. **(O)** Spearman rank correlation analysis of within-group versus between-group pairwise matrix concordance. Obs_T = −0.013, permutation p = 1, indicating no significant difference in matrix concordance within versus between conditions. **(P)** Scatter plot of paired OT-based inter-region similarity values from Control (x axis) versus HIF1A (y axis). Each point represents one region pair. Spearman ρ = 0.978, p < 0.001. Dashed diagonal line indicates identity. **(Q)** Region-level ΔSimilarity (HIF1A − Control) showing minimal per-region changes in mean OT-based inter-region metabolomic coherence across all major brain regions.

To test this, we delivered lentiviral shRNA targeting HIF1α (Lenti-shHIF1α) to the left hippocampus of 5xFAD mice, leaving the right hippocampus as a within-animal internal control. Three months post-injection, social recognition memory was assessed (Figure 5F). Behavioral testing showed that 5xFAD mice receiving Lenti-shHIF1α had significantly improved social memory compared to controls (p = 0.014) (Figure 5G), indicating that local HIF1α knockdown in the hippocampus was associated with improved performance on the social recognition memory assay. Immunofluorescence revealed significant HIF1α reduction on the injected (left) side (p = 0.048) but no significant change on the contralateral (right) side (p = 0.065), indicating that target engagement was restricted to the injection site (Figure 5H). Amyloid-β plaque burden mirrored this asymmetry: plaques were significantly reduced on the injected side (p = 0.017) while the contralateral hemisphere remained comparable to untreated 5xFAD controls, with within-treated left-versus-right comparison confirming a significant injection-side reduction (p = 0.025) (Figure 5I). The absence of contralateral effects on HIF1α expression or amyloid-β load is consistent with the molecular consequences of the intervention being confined to the injected hippocampus, providing within-animal evidence against off-target viral spread. We then examined whether HIF1α knockdown in a single hippocampus altered metabolic coherence across the brain. Coherence was fully preserved (Spearman ρ = 0.978, p < 0.001) (Figure 5J). To assess whether this preservation generalized across the brain, coronal sections were acquired at three matched anatomical depths (depth 1, 2, and 3) capturing three distinct anteroposterior planes and the different anatomical structures contained within each. Inter-regional coherence was preserved across all three depths and at both region and subregion resolution (Supplementary Figures 14, 16, 17, 19, and 20). Across brain regions, multiple TCA cycle intermediates and glycolytic endpoints showed normalization in the differentially regulated metabolite heatmap (Figure 5K). Per-region directional consistency analysis classified the majority of metabolites as predominantly upregulated in the HIF1α cohort (66.0%, n = 97), with 15.0% (n = 22) predominantly downregulated and 19.0% (n = 28) showing mixed responses (Figure 5L). Citric acid, a central TCA cycle intermediate, was significantly normalized in the hippocampal formation and simultaneously across the isocortex, olfactory areas, cortical subplate, thalamus, hypothalamus, and midbrain (p < 0.001 in most regions) (Figure 5M). Per-sample metabolite Log2FC heatmaps showing this coordinated shift across all examined replicate pairs at three anatomical depths are presented in Supplementary Figures 15, 18, and 21. Quantitatively, mean similarity did not differ between control and HIF1α-knockdown brains (p = 0.706, bootstrap Δ = 0.004, 95% CI [−0.028, 0.035]) (Figure 5N). Within-group and between-group similarity structures were statistically indistinguishable (Figure 5O), pairwise similarity values clustered tightly along the identity line (Figure 5P), and region-specific analysis confirmed modest local changes without disruption of global coherence (Figure 5Q). This metabolite normalization occurred in regions that received no direct gene modulation, as the lentiviral transduction and amyloid-β reduction localized to the left hippocampus. Distal metabolite normalization co-occurred with the local intervention even though gene modulation and amyloid reduction localized to the injection site. These data show that local HIF1α knockdown is associated with distal metabolite normalization across regions sharing high metabolic similarity with the hippocampus, consistent with a network-level coordination of brain metabolism.

### Neuronal AOX overexpression in the hippocampus is associated with distal metabolite normalization and improved social memory

To further test whether metabolic coherence functions as a network property of the brain, we pursued a second intervention targeting hippocampal metabolism. Proteomic analysis had revealed perturbed expression of oxidative phosphorylation Complexes II through IV in the 5xFAD brain (Figure 6A), with individual ETC subunit mapping showing both upregulated and downregulated subunits across Complexes II, III, and IV (Figure 6B). Immunostaining of two Complex IV subunits, COX4 and COX5A, in the hippocampus confirmed altered expression in the 5xFAD brain (Figure 6C and 6D), while quantification of COX7a, a structural Complex IV component, showed comparable expression in WT and 5xFAD hippocampus (Supplementary Figure 22A), indicating subunit-selective rather than uniform Complex IV deficits. Based on these findings, we hypothesized that electron flow through the canonical respiratory chain is compromised in the 5xFAD brain. The alternative oxidase (AOX), derived from Ciona intestinalis, is a non-proton-translocating terminal oxidase that accepts electrons from ubiquinol and transfers them directly to molecular oxygen, effectively bypassing Complexes III and IV (Figure 6E).^31,32,33^ Expression of AOX has been shown to restore mitochondrial ubiquinol oxidation in mammalian cells lacking Complex III^31^ and to rescue cytochrome oxidase deficiency phenotypes and mitochondria-mediated neurodegeneration in Drosophila models,^32,33^ establishing AOX as a validated tool for restoring electron flow when the canonical respiratory chain is compromised.^34^ AOX oxidizes the ubiquinol pool upstream of Complex III, thereby reducing electron flux through the Qo site, which represents the dominant source of mitochondrial reactive oxygen species (ROS) in both neurons and astrocytes.^35,36^ Genetic studies in mouse brain have demonstrated that neuron-specific disruption of Complex III triggers severe early ROS-dependent neurodegeneration. This highlighted the unique rescue potential of AOX when ROS is in excess in the central nervous system.^35^

**Figure 6.**
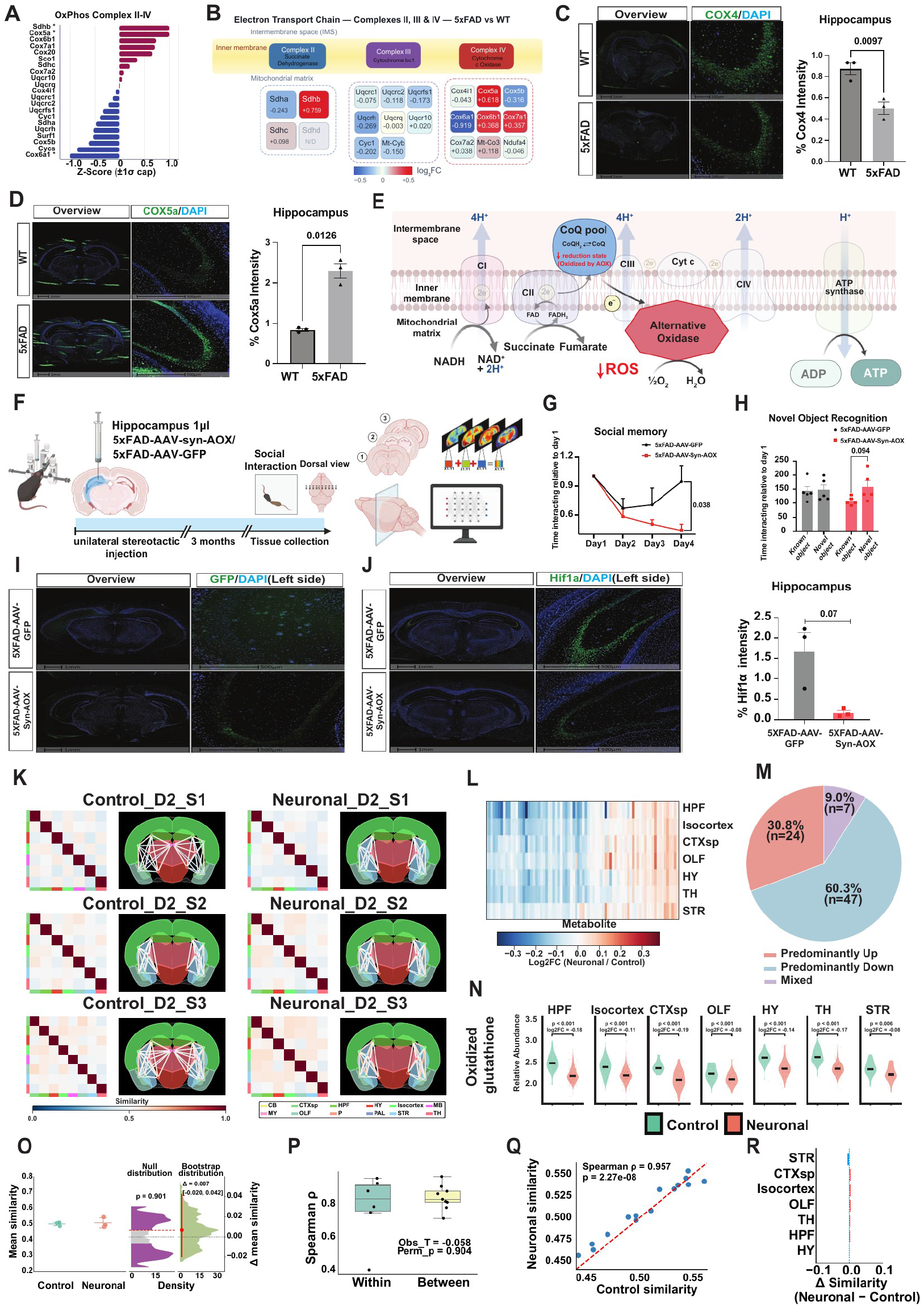
Electron Transport Chain Dysregulation in 5xFAD Brain Is Rescued by Neuronal AOX Expression, Improving Cognitive Function and Reducing HIF-1α. **(A)** Z-score-normalized protein abundance (±1σ cap) for OxPhos complex II--IV subunits in 5xFAD versus WT. **(B)** Schematic of ETC complexes II, III, and IV with individual subunit fold changes (log_2_FC) in 5xFAD versus WT. Blue, downregulated; red, upregulated; gray, not detected. **(C)** Representative coronal sections immunostained for COX4 (green) with DAPI (blue) in WT and 5xFAD hippocampus. Scale bars, 1 mm (overview) and 100 μm (COX4). Right: quantification of % COX4 intensity in hippocampus. n = 3 mice/group; mean ± SEM; [*p = 0*.*0097, two-tailed t-test*. **(D)** Representative coronal sections immunostained for COX5A (green) with DAPI (blue) in WT and 5xFAD hippocampus. Scale bars, 1 mm (overview) and 500 μm (COX5A). Right: quantification of % COX5A intensity in hippocampus. n = 3 mice/group; mean ± SEM; [*p = 0*.*0126, two-tailed t-test*. **(E)** Schematic of alternative oxidase (AOX) bypass in the mitochondrial ETC. AOX accepts electrons from ubiquinol (CoQ), bypassing complexes III and IV to reduce molecular oxygen. **(F)** Schematic of experimental design. 5xFAD mice received unilateral hippocampal injection (1 μL) of AAV-Syn-AOX or AAV-GFP control. Social interaction testing and novel object recognition were performed 3 months post-injection. **(G)** Social memory assessed as interaction time relative to Day 1 across 4 consecutive days. 5xFAD-AAV-GFP (black) and 5xFAD-AAV-Syn-AOX (red). AOX expression rescued social memory habituation. n = 5 mice/group; mean ± SEM; [*p = 0*.*038, two-way ANOVA with multiple comparisons*. **(H)** Novel object recognition test. Time interacting with known versus novel object for 5xFAD-AAV-GFP and 5xFAD-AAV-Syn-AOX groups. n = 5 mice/group; mean ± SEM; [*p = 0*.*094, two-way ANOVA with multiple comparisons*. **(I)** Representative coronal sections showing GFP expression (green) and DAPI (blue) confirming AAV transduction in the hippocampus of 5xFAD-AAV-GFP (top) and 5xFAD-AAV-Syn-AOX (bottom) mice. Scale bars, 1 mm (overview) and 500 μm. **(J)** Representative coronal sections immunostained for HIF-1α (green) and DAPI (blue) in 5xFAD-AAV-GFP (top) and 5xFAD-AAV-Syn-AOX (bottom) mice. Scale bars, 1 mm (overview) and 500 μm. Right: Quantification of % HIF-1α intensity in hippocampus. n = 3 mice/group; mean ± SEM; [*p = 0*.*07, two-tailed t-test*. **(K)** OT-based inter-region metabolomic coherence matrices and corresponding 2D spatial projection maps for three 5xFAD Control (left) and three 5xFAD Neuronal (right) coronal brain sections at anatomical depth 2. n = 3 sections per depth per group. **(L)** Heatmap of per-region metabolite log_2_FC (Neuronal/Control) across brain regions. Rows, brain regions; columns, metabolites. **(M)** Pie chart summarizing the directional consistency of metabolite abundance changes across regions. Metabolites are classified as Predominantly Up (>2/3 of regions increased; 30.8%, n = 24), Predominantly Down (>2/3 of regions decreased; 60.3%, n = 47), or Mixed (neither criterion met; 9.0%, n = 7). **(N)** Region-wise relative abundance of oxidized glutathione in Neuronal (orange) versus Control (green) across brain regions. Oxidized glutathione is consistently downregulated in Neuronal across all examined regions (log_2_FC range: −0.19 to −0.08; p ≤ 0.006 for all regions). **(O)** Gardner-Altman plot comparing mean OT-based inter-region metabolomic coherence between Control (n = 3) and Neuronal (n = 3) samples. Left, individual data points showing sample-level mean similarity per group. Right, bootstrap distribution of the mean difference (Δ = 0.007; 95% CI [−0.020, 0.042]; permutation p = 0.901), indicating no significant global shift in mean OT-based inter-region metabolomic coherence between conditions. **(P)** Spearman rank correlation analysis of within-group versus between-group pairwise matrix concordance. Obs_T = −0.058, permutation p = 0.904, indicating no significant difference in matrix concordance within versus between conditions. **(Q)** Scatter plot of paired OT-based inter-region similarity values from Control (x axis) versus Neuronal (y axis). Each point represents one region pair. Spearman ρ = 0.957, p < 0.001. Dashed diagonal line indicates identity. **(R)** Region-level ΔSimilarity (Neuronal − Control) showing minimal per-region changes in mean OT-based inter-region metabolomic coherence across all major brain regions.

We delivered AAV encoding AOX under the synapsin (SYN) promoter to the left hippocampus of 5xFAD mice, restricting expression to neurons. Control animals received AAV expressing EGFP under the CMV promoter. The experimental design for unilateral hippocampal AOX delivery is shown in Figure 6F. Behavioral testing showed that 5xFAD mice receiving neuronal AOX had significantly improved social memory compared to controls (p = 0.038) (Figure 6G), consistent with the social memory improvement observed with HIF1α knockdown. Novel object recognition performance trended in the same direction toward improvement in AOX-treated mice without reaching conventional significance (p = 0.094) (Figure 6H). GFP immunofluorescence confirmed that transgene expression was confined to the left hippocampus in both groups (Figure 6I). As with the HIF1α intervention, amyloid-β plaque burden was reduced on the injected side only, with no change on the contralateral hemisphere (Supplementary Figure 22B). In line with reduced ROS-driven HIF-1α stabilization, hippocampal HIF-1α immunoreactivity was lower in AAV-Syn-AOX-treated 5xFAD mice than in AAV-GFP controls (Figure 6J). Analysis of metabolic coherence in the neuronal AOX cohort revealed complete preservation of the inter-regional similarity structure (Spearman ρ = 0.957, p < 0.001) (Figure 6K). This preservation of inter-regional coherence was reproduced across additional anatomical depths and at both region and subregion resolution (Supplementary Figures 22C–G, 23, 24, 25, and 26). Coordinated metabolite normalization across multiple regions was evident in the differentially regulated metabolite heatmap (Figure 6L). Per-region directional consistency analysis classified the majority of metabolites as predominantly downregulated in the AOX cohort (60.3%, n = 47), with 30.8% (n = 24) predominantly upregulated and 9.0% (n = 7) showing mixed responses (Figure 6M). Oxidized glutathione, a direct biochemical readout of mitochondrial ROS, was significantly normalized in the AOX cohort across the hippocampal formation, isocortex, cortical subplate, olfactory areas, hypothalamus, thalamus, and striatum (Figure 6N), consistent with AOX-mediated suppression of Complex III-derived ROS. Per-sample metabolite Log2FC heatmaps showing this coordinated shift across all examined replicate pairs at three anatomical depths are presented in Supplementary Figure 27. Quantitatively, mean similarity did not differ between control and AOX-expressing brains (p = 0.901, bootstrap Δ = 0.007, 95% CI [−0.020, 0.042]) (Figure 6O). Within-group and between-group similarity structures did not differ significantly (Figure 6P), pairwise similarity values clustered along the identity line (Figure 6Q), and region-specific analysis revealed the spatial distribution of coherence changes (Figure 6R). Once again, the molecular intervention and amyloid-β reduction localized to the injected hippocampus, while distal metabolite normalization co-occurred in regions sharing high metabolic similarity with the hippocampus, consistent with a network-level character of brain metabolic coordination as captured by coherence.

## Discussion

We define an optimal-transport-derived inter-regional metabolite similarity metric and refer to it as metabolic coherence. Inter-regional molecular similarity has been established at the level of gene expression through large-scale transcriptomic atlases^12,13,14,15,16^ and at the level of neural activity through functional connectivity mapping.^10,11^ Our data are consistent with metabolism being organized at this network level, with metabolites and lipids showing coordinated spatial patterns across anatomically distinct regions. Recent work has begun to extend this framework to metabolism through construction of a human brain metabolic connectome derived from magnetic resonance spectroscopic imaging.^38^ A metabolic connectome of the healthy human brain has now been constructed using whole-brain proton MR spectroscopic imaging, revealing that metabolic similarity forms an organized network topology with a continuous caudal-to-rostral gradient that mirrors structural and cytoarchitectonic organization.^38^ Metabolic coherence differs from these measures because it captures the actual biochemical state of tissue rather than transcriptomic potential or hemodynamic proxy. The astrocyte-neuron lactate shuttle model established that brain metabolism is an actively organized process involving coordinated substrate exchange between cell types,^39^ and our data extend this principle from local cell-cell coupling to inter-regional metabolite similarity at the whole-brain scale. In the framework of network neuroscience, densely connected hub regions incur the highest metabolic costs and are disproportionately vulnerable to pathological attack.^40^ Our observation that the hippocampal formation shows the highest average similarity to other regions in this metric complements these graph-theoretic descriptions of brain organization and suggests that metabolic similarity may reflect shared biochemical inputs across these regions.

The network neurodegeneration hypothesis posits that neurodegenerative diseases target specific intrinsic functional networks, with pathology propagating along the connectivity architecture of the brain.^9^ Cortical hubs correspond spatially to amyloid-β deposition in Alzheimer’s disease.^41^ Metabolic vulnerability propagates through the default mode network in response to distant amyloid pathology.^42^ Intrinsic brain connectivity predicts tau aggregation patterns across all clinical variants of the disease.^43^ Our findings extend this framework into the metabolic domain by showing that metabolic coherence is preserved in 5xFAD brains despite widespread alterations in individual metabolite and lipid levels. The resilience of the coherence structure suggests that the inter-regional organizational scaffold itself is maintained even as individual molecular components shift, consistent with the finding that aerobic glycolysis patterns in young adults predict both amyloid vulnerability and cognitive resilience in later life.^2,3,37^ The inter-regional similarity structure is reproducible in 5xFAD mice despite metabolite-level alterations. Individual metabolites in 5xFAD brains shift in a coordinated fashion across regions, preserving the relative inter-regional pattern even as absolute levels change. Several non-mutually-exclusive mechanisms could produce this coordinated shifting: cell-autonomous responses to a shared systemic stimulus such as circulating metabolites or blood-borne ROS, trans-regional signaling through long-range axonal projections, glial network coupling via the astrocytic syncytium, or redistribution of substrates through the cerebrovasculature. The current data do not distinguish among these candidates, but the observation that the hippocampal formation shows the highest average coherence with other regions and that local hippocampal intervention is associated with distal effects is most consistent with mechanisms that involve the hippocampus as a coordinating hub rather than a passive recipient of a shared external input. Recent work has demonstrated that preservation of youthful aerobic glycolysis patterns is associated with cognitive resilience in amyloid-positive individuals^37^. Whether metabolic coherence contributes to this resilience is an open question.

The central finding of our study is that two mechanistically distinct interventions, each confined to the left hippocampus, were associated with distal metabolite normalization and improved social memory in 5xFAD mice. HIF1α knockdown targets a transcriptional master regulator of glycolysis,^30^ while AOX overexpression provides an alternative electron acceptor that bypasses Complex III and IV of the mitochondrial electron transport chain.^31^ Despite engaging entirely different metabolic pathways, both interventions were associated with improved social memory and with metabolite normalization in regions sharing high metabolic similarity with the hippocampus, including the isocortex, thalamus, hypothalamus, and olfactory areas. Two distinct metabolic interventions in the same region were associated with overlapping distal metabolite changes. Both interventions improved performance on the social recognition memory assay. HIF1α knockdown (p = 0.014) and neuronal AOX (p = 0.038) both reached conventional significance, and the difference in effect size may reflect the distinct positions of these interventions along the ETC-ROS-HIF1α signaling axis. AOX acts upstream by eliminating the ROS source at the Qo site, whereas HIF1α knockdown acts downstream on the transcription factor itself, potentially leaving residual ROS-mediated damage uncorrected. Alternatively, differences in viral vector kinetics (lentiviral versus AAV delivery) or the dual pro-survival and pro-death roles of HIF1α in neurons may contribute to the differential behavioral response. Our findings provide a mechanistic complement to the observation that amyloid-β in distant brain regions induces regional hypometabolism through functional connectivity,^42^ as tau burden in highly connected nodes progressively weakens their functional connectivity.^44^ Those studies show that pathology propagates through networks to impair metabolism. The contralateral hippocampus served as a within-animal control for both interventions: Lenti-shHIF1α produced no significant change in HIF1α expression (p = 0.065) or amyloid-β load on the uninjected side (Figure 5H, 5I), and AAV-Syn-AOX expression was restricted to the injected hippocampus by GFP staining (Figure 6I) with amyloid-β reduction confined to that side (Supplementary Figure 22B). This spatial restriction at the injected hemisphere argues against off-target viral spread to distal regions millimeters to centimeters away and supports the interpretation that distal metabolite normalization reflects network-level metabolic coordination rather than direct viral transduction at distal sites. These data identify a reproducible inter-regional metabolomic similarity structure in the mouse brain. In 5xFAD mice, this structure is preserved despite metabolite-level alterations. Local hippocampal metabolic interventions are associated with metabolite shifts in distal regions, particularly regions metabolically similar to the hippocampus, suggesting that metabolic coherence may reflect a network-level organization of brain metabolism. Neuronal AOX expression under the synapsin promoter was associated with both behavioral improvement and distal metabolite normalization in 5xFAD mice.

Several aspects of this study open important avenues for future investigation. Future studies using isotope tracing or genetically encoded metabolic sensors could determine whether metabolites are physically transported between coherent regions or whether coherence arises from shared regulatory programs acting independently in each area. Neuronal AOX expression in the hippocampus was associated with distal metabolite normalization. Independent support comes from recent spatial metabolomics work showing that focal ischemic injury produces sustained metabolic changes in the histologically unaffected ipsilateral cortex,^45^ indicating that local metabolic perturbations can co-occur with metabolite changes in distant brain regions. Future work using cell-type-restricted interventions, including astrocytic AOX, will be needed to dissect the cellular basis of this association. MALDI-MSI provides a snapshot of the metabolic landscape at the time of tissue collection, and emerging live-imaging metabolomics approaches will be essential for resolving the temporal dynamics of metabolic coordination and the kinetics of distal metabolite changes. Determining whether metabolic coherence is conserved in the human brain is an important next step, and the rapid maturation of spatial metabolomics technologies for human tissue makes direct validation in human postmortem and surgical specimens increasingly tractable.

## Acknowledgments

This study was supported by National Institutes of Health (NIH) grants R01AG066653, R01CA266004, R01AG078702, R01CA288696, and RM1NS133593 to R.C.S., R35NS116824 to M.S.G., and R35GM142701 to L.C. Z.L. is supported by the MBI Gator NeuroScholar Program. T.M. is supported by NIH T32 HL134621. C.S. is supported by the NMPT program under NIH T32 HD043730. The UF Neuromedicine Human Brain and Tissue Bank is supported by the McKnight Brain Institute, the Center for Translational Research in Neurodegenerative Diseases at the University of Florida, and NIH grant P30AG066506. Large language models (Claude) were used for grammar checking and proofreading.

## Author Contributions

Conceptualization, R.C.S.; Methodology, R.C.S., X.M., and L.C.; Formal Analysis, X.M., Z.L., and L.C.; Investigation, Z.L., X.M., R.A.R., T.M., S.Q., J.S., L.W., A.M.R., C.S., B.Z., V.B-C., N.R., K.W., A.T., and R.L.; Resources, C.W.V.K., M.S.G., and N.S.C.; Writing – Original Draft, R.C.S.; Writing – Review & Editing, R.C.S., N.S.C., C.W.V.K., M.S.G., and L.C.; Funding Acquisition, R.C.S., M.S.G., and L.C.; Supervision, R.C.S., M.S.G., and L.C.

## Methods

### Animals

5xFAD transgenic mice (JAX stock #034848) and their wild-type (WT) littermates were obtained from The Jackson Laboratory. All animals were housed under standard laboratory conditions and maintained on a 12-hour light/dark cycle with ad libitum access to food and water. Stereotaxic intracranial injection procedures were performed at approximately 3 months of age. Behavioral testing and pathological examinations were conducted when animals reached 6 to 8 months of age. All animal protocols were approved by the University of Florida Institutional Animal Care and Use Committee (IACUC).

### Stereotaxic injections

All surgeries were performed under stereotaxic guidance. For HIF1α knockdown experiments, lentivirus encoding shRNA targeting HIF1α (TL517255V; OriGene) or control scrambled shRNA lentivirus (TR30023; OriGene) was injected unilaterally into the left hippocampus of 5xFAD mice (AP: −0.56 mm, ML: 1.1 mm, DV: −2.2 mm from bregma). Lentiviral particles were delivered in a 1 µL volume at a constant rate of 0.5 µL/min using a Hamilton syringe. Following infusion, the injection needle was left in place for 5 minutes to allow adequate viral diffusion and to prevent backflow before being slowly withdrawn.

For alternative oxidase (AOX) overexpression experiments, mice received unilateral stereotaxic injections of either rAAV-Syn-AOX-GFP (VB250331-1441kha; VectorBuilder) for neuron-specific expression or AAV-CMV-AOX-GFP (VB250331-1443mxu; VectorBuilder) for general cellular expression into the left hippocampus using the same coordinates. Adeno-associated viral particles were delivered in a 1 µL volume at an injection rate of 0.5 µL/min. The syringe was maintained in place for 5 minutes post-injection to ensure diffusion before removal. Control animals received AAV expressing EGFP under the CMV promoter. To ensure sufficient viral expression, all behavioral assays were conducted 3 months post-injection.

### Sample collection and preparation

Fresh-frozen brain tissues were collected according to previously established protocols. Mice were rapidly euthanized by cervical dislocation, and brains were immediately excised. To eliminate residual blood, brain tissues were rinsed once in phosphate-buffered saline (PBS) and subsequently washed twice with deionized water. Whole brains were then snap-frozen over liquid nitrogen to preserve molecular integrity and stored at −80°C until further processing.

### MALDI mass spectrometry imaging

Frozen brain tissues were cryosectioned at −20 °C (Leica CM1860) at a thickness of 10 µm. Coronal sections were thaw-mounted onto Bruker IntelliSlides, vacuum-sealed, and stored at −80 °C until analysis. Immediately prior to matrix application, slides were brought to room temperature and dried via vacuum desiccation for 1 hour to ensure surface dehydration and metabolite stability. Spatial metabolomics and lipidomics were performed sequentially on the same tissue sections. The matrix N-(1-Naphthyl) ethylenediamine dihydrochloride (NEDC) was applied using an HTX M5 TM-Sprayer (HTX Imaging). A 7 mg/mL NEDC solution in 70% methanol was delivered using the following optimized parameters: 14 passes, 30 °C nozzle temperature, 0.06 mL/min flow rate, 10 psi nitrogen pressure, and a 50 °C tray temperature.

### MALDI-timsTOF imaging acquisition

Data were acquired on a timsTOF fleX mass spectrometer (Bruker Daltonics) equipped with a 10 kHz SmartBeam 3D laser. To maximize reproducibility and minimize user bias, region definition and instrument setup were automated using the autopilot feature in flexImaging v6.0. High-resolution optical scans (.tif) were used to generate a standardized geometry (flexImaging.mis) for consistent region masking and pixel alignment across all omics runs. Imaging was performed in negative ion mode using a 46 µm laser raster to achieve a 50 µm pixel pitch.

Metabolomics. MS1 spectra were collected from m/z 20–750 at 80% laser power (1 burst of 396 shots). Global instrument settings included a 30 V MALDI plate offset, −60 V deflection 1 delta, and 200 Vpp for Funnels 1 and 2 and the Multipole RF. Collision energy was set to 7 eV with a 700 Vpp collision RF.

Lipidomics. Following metabolomics acquisition, lipidomics data were immediately collected from the same pixels (m/z 300–2000) at 80% laser power (1 burst of 300 shots), utilizing the same raster and transfer parameters to ensure perfect spatial registration between the two datasets.

### MALDI raw data alignment and normalization

Post-acquisition, raw data were imported into SCiLS Lab (v2024b, Bruker Daltonics) for spectral alignment and normalization. Spatially resolved ion distributions were extracted, and total ion count (TIC) normalization was applied to account for matrix heterogeneity and tissue effects. The aligned and normalized ion intensity matrices were then exported and used as input for the downstream metabolomic coherence analyses described below.

Replication. Three-dimensional sagittal MALDI-MSI volumes were generated from one wild-type and one 5xFAD brain (n = 1 per genotype). Coronal cohorts comprised three independent biological replicates per group: n = 3 wild-type and n = 3 5xFAD for genotype comparison; n = 3 5xFAD-Lenti-shHIF1α and n = 3 5xFAD-Lenti-shScrambled control for the HIF1α intervention; and n = 3 5xFAD-AAV-Syn-AOX and n = 3 5xFAD-AAV-CMV-GFP control for the neuronal AOX intervention. Within each intervention cohort, three coronal sections per replicate were acquired at three matched anatomical depths (depth 1, 2, and 3), corresponding to three distinct anteroposterior planes that capture different anatomical structures along the rostral-caudal axis (depth 1, anterior; depth 2, mid; depth 3, posterior), and all between-group analyses were performed independently within each depth stratum.

### Behavioral testing

#### Social memory assay

Social recognition memory was assessed using a previously published paradigm. On the day of testing, experimental mice were acclimated to the behavioral room for 30 minutes before being individually placed in a clean cage for 10 minutes of free exploration. An unfamiliar target mouse was subsequently introduced for a 10-minute interaction phase. This protocol was repeated daily across five consecutive days, with the experimental subject encountering the same target mouse during each trial. Behavioral sessions were continuously recorded and objectively analyzed using the deep-learning pose estimation software DeepLabCut.

#### Novel object recognition

Novel object recognition memory was assessed using a two-phase paradigm. Following a 30-minute room acclimation, mice underwent a 10-minute familiarization trial in which they were placed in an open-field chamber containing two identical objects for free exploration. One hour later, a 5-minute recognition trial was conducted in which one of the familiar objects was replaced with a novel object of identical dimensions but entirely different shape and visual characteristics. Behavioral sessions across both phases were continuously recorded and quantified using the ANY-maze (Stoelting Co.) tracking system.

#### Open field test

The open field test was conducted to measure basal locomotor activity. Prior to assessment, animals were transferred to the testing room and allowed a 30-minute acclimation period. Each mouse was individually placed into an open-field chamber (40 × 40 × 40 cm) and allowed to explore freely for 30 minutes. All behavioral metrics, including total distance traveled and duration spent in the center area, were continuously tracked and analyzed using ANY-maze (Stoelting Co.) video tracking software.

### Immunofluorescence and immunohistochemistry

Immunostaining was performed on fresh-frozen brain specimens. Coronal sections (10 µm) were obtained using a Leica cryostat (CM1950, Leica Microsystems, Germany). Cryosections were briefly post-fixed in 1% paraformaldehyde (PFA) solution to preserve tissue architecture. Following fixation, slides were systematically rehydrated and subjected to antigen retrieval to unmask immunoreactive epitopes. To minimize non-specific antibody binding and facilitate cellular penetration, sections were permeabilized with 0.5% Triton X-100 and blocked using 10% normal goat serum for 1 hour at room temperature.

Processed tissues were incubated overnight at 4°C with the following primary antibodies: GFAP (astrocyte marker; 1:100, GTX85454; GeneTex), 6E10 (amyloid-β plaque marker; 1:100, 803015; BioLegend), HIF1α (1:100, GTX127309; GeneTex), and GFP (1:200, GTX113617; GeneTex). Following three washes with PBS to remove unbound antibodies, sections were incubated with corresponding fluorophore-conjugated secondary antibodies for 1 hour at room temperature, protected from light. Digital whole-slide images were acquired using a VS200 scanner. Regions of interest were selected and quantitative image analysis was performed using HALO software (Indica Labs).

### Proteomics

Brain tissue samples were resuspended in high-salt lysis buffer (2.0 M NaCl, 5.0 mM EDTA, pH 7.4, supplemented with 1× protease inhibitor cocktail; 1.0 mL total volume). Mechanical disruption was achieved by repeated passage through a 20-gauge needle (10 strokes) followed by probe sonication. The homogenate was subjected to ultracentrifugation at 45,000 rpm for 15 minutes at 4°C, after which the resulting supernatant was discarded. The pellet was resuspended in sodium carbonate buffer (0.1 M Na_2_CO_3_, 1.0 mM EDTA, pH 11.3; 1.0 mL) and incubated on ice for 30 minutes prior to a second ultracentrifugation step (45,000 rpm, 45 minutes, 4°C). Following centrifugation, the supernatant was removed, and the membrane-enriched pellet was solubilized in urea lysis buffer (8.0 M urea, 1.0 M NaCl, 4% w/v CHAPS, 100 mM DTT, 200 mM Tris-HCl, pH 8.0; 1.0 mL) with heating at 50°C for 45 minutes to facilitate protein denaturation.

Alkylation of free thiol groups was carried out by adding iodoacetamide (IAA; 55 mg) and incubating at room temperature in the dark for 45 minutes. Proteins were then precipitated by sequential addition of chloroform (1.5 mL), methanol (4.5 mL), and water (5.0 mL), and recovered by centrifugation at 3,000 × g for 5 minutes. The protein pellet was washed once with methanol, re-centrifuged, and reconstituted in 50 mM ammonium bicarbonate buffer (pH 8.0; 500 µL). Total protein concentration was determined by BCA assay. For proteolytic digestion, 100 µg of membrane protein was incubated with sequencing-grade trypsin at an enzyme-to-substrate ratio of 1:20 (w/w) at 37°C for 24 hours.

Tryptic peptides were analyzed on an Ultimate 3000 RSLCnano system coupled to a Thermo Eclipse Tribrid mass spectrometer. Chromatographic separation was performed using a nano-LC column (15 cm length, 75 µm internal diameter) packed with 3 µm C18 reverse-phase material. Peptides were eluted over a 180-minute gradient of increasing acetonitrile concentration in 0.1% formic acid. Precursor ion scans were acquired at 120,000 resolution in the Orbitrap analyzer. Precursor ions within a 3-second cycle time were selected for fragmentation by higher-energy collisional dissociation (HCD). Charge state screening was enabled, and precursors with unknown or +1 charge states were excluded. Dynamic exclusion was set to 30 seconds. Fragment ions were analyzed in the Orbitrap at 30,000 resolution.

Raw data were processed using Proteome Discoverer (v3.0, Thermo Fisher Scientific). Peptide identification was performed using the CHIMERYS search algorithm with a summed posterior error probability (Sum PEP) threshold of 15. Precursor and fragment mass tolerances were set to 5 ppm and 10 ppm, respectively. Variable modifications included deamidation of asparagine and glutamine, carboxymethylation of cysteine, and oxidation of methionine. Differential expression analysis and pathway enrichment for oxidative phosphorylation, glycolysis, and TCA cycle pathways were performed on the identified protein abundances.

### Data Preprocessing

MALDI imaging data were preprocessed before analysis. Specifically, total ion current (TIC) normalization was applied to account for differences in overall signal intensity across pixels, followed by log-transformation using *log*(1 + *x*). A scaling factor of 10000 was applied to stabilize numerical ranges and improve comparability across samples. The processed data were used for subsequent analysis, including the generation of the PC1 image for use as the fixed image in registration, as well as for computing metabolomic coherence and other downstream metrics.

For three-dimensional data consisting of serial sections, inter-section alignment was performed in advance using the MetaVision3D framework. This step produced an aligned MALDI volume, ensuring spatial consistency across sections prior to atlas registration.

### MALDI Atlas Registration

We provided both manual and automatic strategies for registering Allen Brain Atlas and MALDI brain images. Manual registration was used when sufficient anatomical prior knowledge was available and high registration accuracy was required, whereas automatic registration offered a more efficient alternative when prior information was limited or when approximate alignment was sufficient for downstream analysis.

#### Manual registration using the QUINT workflow

Manual registration between MALDI brain images and the Allen Brain Atlas was performed using the QUINT workflow, which consists of two major modules, QuickNII and VisuAlign. This workflow combines global anatomical alignment and local deformation correction, enabling atlas annotations to be accurately mapped onto MALDI sections.

#### Registration preprocessing

To preserve the integrity of the original MALDI data, the first principal component (PC1) across all metabolites based on preprocessed data was used as the fixed image, while the atlas image from the QUINT package (Allen Brain Atlas, CCFv3, 25 µm resolution) was treated as the moving image. The use of PC1 is motivated by its ability to capture the dominant spatial variation in metabolite intensities, producing a smooth, high-contrast representation of tissue morphology that facilitates robust cross-modality alignment. Using the atlas as the moving image avoids resampling or distorting the original MALDI measurements.

#### QuickNII: global alignment

The first step of manual registration was performed in QuickNII to establish global correspondence between each MALDI section and the Allen Brain Atlas. The PC1 image was imported as the target section, and the atlas volume from the QUINT package was used as the reference. For each section, the corresponding atlas plane was first identified by visually matching major anatomical landmarks, such as the overall brain outline, ventricular structures, cortical boundaries, hippocampal shape, and other prominent regional features visible in the PC1 image.

After selecting the most appropriate atlas plane, the section was manually adjusted in QuickNII through translation, rotation, and scaling to achieve coarse alignment between the atlas and MALDI image. This step focused on matching the global orientation, section position, and overall morphology rather than fine regional details. The purpose of QuickNII was to provide a reliable initialization for subsequent refinement by ensuring that the atlas slice approximately overlapped the MALDI tissue section in anatomical space.

For serial sections, the alignment was performed sequentially to maintain consistency across neighboring slices. The estimated section position and orientation from adjacent slices were used as references when available, which helped preserve smooth anatomical progression throughout the series.

#### VisuAlign: local refinement

Following global alignment in QuickNII, the resulting atlas-to-section mapping was further refined in VisuAlign to correct local mismatches that could not be captured by rigid or affine transformations alone. These mismatches may arise from tissue deformation during sample preparation, sectioning artifacts, differences in imaging contrast between MALDI and atlas images, and local non-linear shape variations.

In VisuAlign, the coarse alignment obtained from QuickNII was loaded as the initial registration. Corresponding landmarks were then manually placed between the atlas and MALDI image, with emphasis on anatomically meaningful boundaries and internal structures that were clearly identifiable in both modalities. Based on these landmarks, non-linear warping was applied to deform the atlas locally so that regional contours more closely matched the MALDI-derived tissue morphology.

The refinement process was performed iteratively. After each adjustment, the overlaid atlas boundaries were visually inspected and further corrected when necessary, particularly in regions with complex geometry or higher local distortion. This procedure enabled the atlas annotations to follow the actual tissue shape more closely while preserving global anatomical plausibility.

### Automatic registration using customized pipeline

Automatic registration between the MALDI imaging and the Allen Brain Atlas was performed using a customized multi-stage pipeline implemented in Python. This pipeline was designed progressively align the two modalities through similar two steps: global alignment and local refinement.

#### Registration preprocessing

Similar to manual registration, the fixed images MALDI PC1 volume (NIfTI format) and the moving images Allen Brain Atlas (TIFF format) were loaded using SimpleITK (sitk.ReadImage) and tifffile.imread, respectively. To ensure consistency across modalities, both MALDI and atlas volumes were further adjusted to share a common orientation and coordinate system prior to registration.

To improve robustness for multimodal registration, intensity normalization was applied using a percentile-based scaling approach implemented with NumPy, which reduces the influence of outliers and stabilizes intensity distributions. In addition, histogram matching (sitk.HistogramMatchingImageFilter) was performed to further reduce intensity discrepancies between modalities.

#### Global alignment

Global alignment was performed to establish coarse correspondence between the atlas and MALDI volumes. To account for differences in anatomical depth and slicing schemes, an anchor-based slice mapping strategy was applied.

Corresponding slices between the atlas and MALDI volumes were manually specified, and intermediate slices were interpolated to establish a consistent slice-to-slice mapping. The atlas volume was then padded and spatially adjusted to match the MALDI field of view.

A rigid transformation was first estimated using sitk.ImageRegistrationMethod, with Mattes Mutual Information as the similarity metric and initialization via sitk.CenteredTransformInitializer, correcting for global translation and rotation differences.

This was followed by affine registration using sitk.AffineTransform, which further accounts for global scaling and shearing effects between the two modalities.

To capture non-linear anatomical differences at the whole-brain level, a deformable registration step was subsequently applied using a B-spline transformation initialized with sitk.BSplineTransformInitializer. A multi-resolution optimization strategy (coarse-to-fine pyramid) was employed to improve convergence and robustness, allowing the model to progressively capture deformation from large-scale structures to finer anatomical details.

#### Local refinement

To further improve alignment in anatomically complex regions where global transformations are insufficient, local refinement was performed on top of the global transformation. Regions of interest were extracted using sitk.RegionOfInterest, focusing on areas with higher structural variability or residual misalignment. Within each region, additional B-spline-based deformable registration was performed to refine local correspondence between atlas and MALDI images. These local transformations were converted into displacement fields using sitk.TransformToDisplacementField and integrated into the global deformation field using sitk.Paste. To ensure spatial smoothness and avoid discontinuities at region boundaries, the combined displacement field was regularized using Gaussian smoothing (sitk.DiscreteGaussian). Finally, all transformations (rigid, affine, and deformable) were combined using sitk.CompositeTransform, and the final transformation was applied to warp the atlas into MALDI space using sitk.Resample. The registered atlas volume and corresponding deformation fields were saved using sitk.WriteImage for downstream analysis.

#### Generation of registered atlas annotations

After either manual or automatic registration, the final transformation was used to project atlas annotations from the Allen Brain Atlas into the MALDI image space. Each MALDI pixel was then assigned to an anatomical region according to the transformed atlas labels.

We defined two levels of anatomical granularity based on the atlas annotation hierarchy. At a coarse level, regions were grouped into 12 major anatomical sets derived from the Allen Brain Atlas structure sets. There sets represent high-level brain divisions. At a finer level, we retained the original subregion annotations within each structure set. For clarity and consistency, we refer to the coarse anatomical structure sets as regions, and the finer anatomical units as subregions. This hierarchical organization enables flexible analysis across scales, allowing both region-level and subregion-level investigation.

### Metabolomics Coherence Analysis

After atlas registration, anatomical annotations were projected into the MALDI image space, allowing each MALDI pixel to be assigned to a corresponding anatomical label. Based on these registered annotations, metabolite intensity profiles were extracted for all pixels within each annotated anatomical unit. As described above, we used two levels of anatomical granularity in downstream analysis: a coarse level consisting of 12 atlas-derived regions and a finer level consisting of subregions nested within each region.

#### Preprocessing and Quality Control

For each sample, the MALDI data were stored in an AnnData object after quality control and preprocessing. Pixels belonging to the target sample were first subsetted, and the corresponding metabolite intensity matrix was combined with pixel-level metadata, including spatial coordinates and anatomical annotations. For region-level analysis, the original atlas labels were mapped to the 12 coarse anatomical regions using a predefined region-to-set dictionary. This enabled the same computational framework to be applied at either the region or subregion level by changing the annotation granularity.

Before pairwise comparison, anatomical units with insufficient valid pixels were excluded to improve robustness. Pixels with non-zero metabolite intensity were considered valid, and anatomical units were retained only if they contained a sufficient number of valid pixels. The minimum pixel threshold was adaptively determined based on data dimensionality (2D vs. 3D) and the level of anatomical granularity (region vs. subregion), ensuring stable estimation of metabolite distributions while preserving adequate coverage across units.

#### Optimal transport-based distance between anatomical units

To quantify metabolomic similarity between anatomical regions, we modeled each region as a probability distribution over pixel-level metabolite profiles and computed pairwise inter-region distances using an optimal transport (OT) framework.

Prior to distance computation, pixels with non-finite values or all-zero intensity vectors were excluded to remove background and artifact-dominated measurements. The remaining pixels were treated as empirical samples from the region-specific metabolomic distribution.

For a given region *r*_*i*_, let 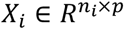 denote the metabolite intensity matrix, where *n*_*i*_ is the number of pixels in the region and *p* is the number of measured metabolites. Each row of *X*_*i*_ corresponds to the full metabolite profile of a single pixel, representing a point in the p-dimensional metabolite space. Each region *r*_*i*_ was then represented as a weighted empirical measure

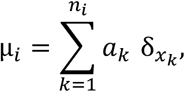

where *x*_*k*_ ∈ *R*^*p*^ denotes the metabolite vector of the k-th pixel and 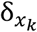 is the Dirac measure placing unit mass at *x*_*k*_. We adopted a non-uniform weighting scheme to account for the heterogeneous signal distribution inherent in MALDI imaging data. Specifically, each pixel was assigned a weight proportional to its total metabolite intensity across all features:

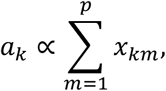

followed by normalization such that 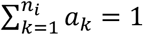.

This formulation down-weights low-intensity or noisy pixels while allowing high-signal pixels to contribute more prominently to the regional distribution since high-signal pixels are more likely to reflect biologically meaningful molecular content.

Pairwise distances between regions were computed using the entropy-regularized optimal transport (Sinkhorn) formulation, as implemented in the GeomLoss package. Given empirical measures µ_i_ and µ_*_, the Sinkhorn distance is defined as

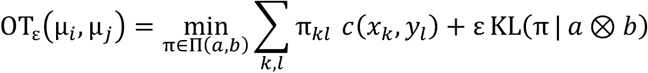

where:

1. *k* = 1, …, *n*_*i*_ and *l* = 1, …, *n_j_* index pixels in regions *i* and *j*;
2. 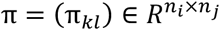 is the transport plan, where *π_kl_* represents the amount of mass transported from pixel *x*_*k*_ to *y_l_*;
3. Π(*a, b*) denotes the set of admissible transport plans satisfying marginal constraints:

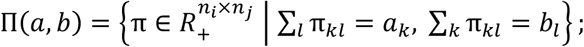

*c*(*x*_*k*_, *y*_l_) = |*x*_*k*_ – *y*_l_|^62^ is the squared Euclidean ground cost measuring dissimilarity between metabolite profiles;

1. KL(⋅) denotes the Kullback–Leibler divergence;
2. ε > 0 is the entropic regularization parameter controlling the smoothness of the transport plan.

In our implementation, we used exponent *p* = 2 and blur parameter σ = 0.1, corresponding to a regularization strength of ε = σ^p^ = 0.01. This relatively small regularization ensures that the solution remains close to the exact Wasserstein distance while enabling efficient computation via Sinkhorn iterations. The resulting pairwise OT distance matrix was used as the region-to-region metabolomic distance measure.

#### Pairwise coherence matrix construction

For each sample, OT distances were computed for all pairs of retained anatomical units, resulting in a symmetric region-by-region distance matrix. Let *D_ij_* denote the OT distance between anatomical units *i* and *j*. To improve numerical stability and compress the scale of large values, distances were transformed using 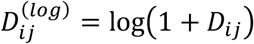. These transformed distances were then converted into a similarity measure, which we refer to as metabolomic coherence, using a Gaussian kernel:

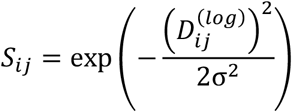

Here, σ was defined as the median of the upper-triangular entries of the log-transformed distance matrix within each sample. This adaptive bandwidth selection normalizes the similarity scale according to the overall distance structure of the sample.

The resulting matrix *S* is a symmetric coherence matrix in which larger values indicate greater similarity between the metabolite distributions of two anatomical units.

#### Correlation-based metabolomic coherence

As a complementary and computationally efficient alternative to optimal transport, we quantified inter-region metabolomic coherence using correlation-based measures. For each sample, MALDI pixel-level data were first grouped by anatomical region. To ensure robustness, pixels were filtered based on feature prevalence. Specifically, for each pixel, the proportion of metabolites with non-zero intensity was computed, and only pixels exceeding a predefined prevalence threshold were retained. Regions with insufficient numbers of valid pixels were excluded from further analysis. In addition, a consistent set of regions shared across samples was used to ensure comparability.

For each retained region, a representative metabolite profile was obtained by aggregating pixel-level measurements. Specifically, the mean of metabolite intensities was computed across all pixels within each region. Let 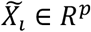 denote the aggregated metabolite vector for region *i*, where *p* is the number of metabolites. To remove scale differences across metabolites, feature-wise standardization was applied across regions. For each metabolite, values were transformed to have zero mean and unit variance across all regions. Pairwise similarity between regions was then computed using Pearson correlation. Specifically, for regions *i* and *j*, the correlation was defined as 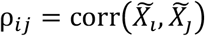, where corr(⋅,⋅) denotes the Pearson correlation coefficient. This resulted in a symmetric region-by-region correlation matrix, where higher values indicate more similar metabolite profiles.

This correlation-based approach captures similarity in average metabolite patterns across regions and serves as a baseline method for comparison with distribution-based measures such as optimal transport.

### Statistical Analysis

#### Two-sample comparison using 3D data

Due to substantial experimental cost and acquisition complexity associated with 3D MALDI imaing, each condition (WT and 5xFAD) was represented by a single 3D MALDI sample in this study. As a result, only one inter-region similarity matrix is available per condition. Under this setting, we performed two-sample comparisons at both the matrix and distribution level to assess differences in inter-region metabolomic similarity, capturing two complementary aspects: pattern concordance and similarity strength.

#### Matrix-level concordance analysis via permutation testing

To assess whether the overall pattern of inter-region relationships is preserved across conditions, we performed a matrix-level concordance analysis that evaluates whether region pairs that are highly similar in one condition remain highly similar in the other. This analysis is designed for settings in which only a single similarity matrix is available per condition (i.e., no biological replicates). For each condition, a symmetric region-by-region similarity matrix was constructed using optimal transport. To avoid redundancy, only the upper triangular entries (excluding the diagonal) were extracted from each matrix, resulting in a vector representation of all unique region pairs.

The concordance between two matrices was quantified using the Spearman rank correlation coefficient:

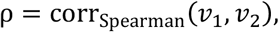

where *v*_1_ and *v*_2_ denote the vectorized upper-triangular entries of the two matrices. This measure captures the consistency in relative similarity patterns across region pairs, independent of absolute scale differences.

To assess statistical significance, we performed a permutation test based on node-label randomization. Specifically, for each permutation, the region labels of one matrix were randomly permuted simultaneously along rows and columns, preserving matrix symmetry while disrupting the correspondence between regions. The Spearman correlation between the permuted matrix and the original matrix from the other condition was then recomputed. Repeating this procedure *N* = 1000 times generated a null distribution of correlation coefficients under random label assignment. The permutation p-value was calculated as 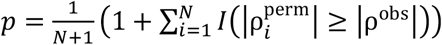, where ρ^obs^ is the observed correlation and 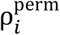 are the permuted correlations.

This analysis evaluates whether the observed similarity pattern between regions is significantly preserved across conditions, beyond what would be expected by chance.

#### Distribution-level comparison of inter-region similarity

Complementing the pattern-level analysis, we further assessed whether the overall strength of inter-region metabolomic similarity differs across conditions using a distribution-level analysis that compares the magnitude of similarity values between two groups. Similarly, this approach is intended for scenarios with a single similarity matrix per condition. Only the upper triangular entries of each similarity matrix were used for analysis. To visualize the distribution, we used histogram and empirical cumulative distribution function (ECDF) to visualize the distribution.

To formally test whether the similarity distribution differ between conditions, we applied the two-sample Kolmogorov-Smirnov (KS) statistic, defined as 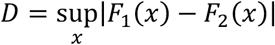, where *F*_1_ and *F*_2_ denote the empirical cumulative distribution functions of the two groups. The KS statistic captures the maximum difference between the two distributions. To assess statistical significance, we performed a permutation-based KS test. Specifically, similarity values from both conditions were pooled and randomly reassigned into two groups while preserving the original sample sizes. For each permutation, the KS statistic was recomputed to generate a null distribution under the hypothesis of no group difference. This procedure was repeated *N* = 1000 times. The permutation p-value was calculated as 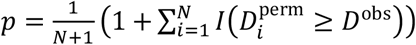, where *D*^obs^ is the observed KS statistic and 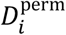 are the permuted values.

This analysis evaluates whether the overall distribution of inter-region metabolomic similarity differs significantly between conditions, providing a complementary perspective to matrix-level concordance analysis.

#### Direction stability Analysis

To address the limited availability of 3D MALDI data, we investigated whether 2D sections can serve as a reliable approximation of 3D spatial organization. Specifically, we hypothesized that if metabolite spatial distributions are stable across anatomical axes, then 2D measurements can capture the essential spatial structure observed in 3D data. To this end, we performed a direction stability analysis based on an equivalence testing framework to quantify the extent of variation in metabolite distributions across anatomical directions.

For each metabolite, spatial variation was evaluated independently along each anatomical axis (x, y, and z). Within a given axis, pixels were grouped into slices, and only slices containing at least 100 pixels were retained to ensure reliable estimation. To control for sampling imbalance across slices, a maximum 4000 pixels per slice was randomly subsampled when necessary. The global distribution *G* was defined as the pooled distribution across all slices. To improve computational efficiency, *G* was subsampled to at most 10,000 observations when necessary. To quantify the deviation of individual slices from the global distribution, we computed the Wasserstein distance between each slice and *G*, and defined the test statistic as the average slice-to-global distance:

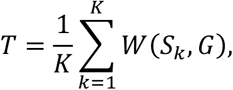

where *W*(⋅,⋅) denotes the one-dimensional Wasserstein distance.

To determine whether the observed variation is meaningful, we estimated a metabolite-specific baseline variability δ using a split-half resampling strategy within slices. For each slice, pixel intensities were randomly partitioned into two disjoint subsets of equal size, and the Wasserstein distance between the two subsets was computed. This procedure was repeated 50 times per slice, and the baseline variability δ was defined as the 95th percentile of all resulting within-slice distances. Here, δ represents the intrinsic variability (or noise level) of the metabolite distribution within slices, capturing the degree of variation expected due to sampling variability and measurement noise rather than true spatial differences. As such, δ serves as a reference scale against which between-slice variation is evaluated for identifying directionally stable metabolites.

To quantify uncertainty in the test statistic, we constructed confidence intervals for *T* using bootstrap resampling. Within each slice, pixel intensities were resampled with replacement, and the statistic *T* was recomputed for each bootstrap replicate. This procedure was repeated 500 times, and a 95% confidence interval [CI_low_,CI_high_] was obtained from the empirical distribution.

A metabolite was considered directionally stable along a given axis if the upper bound of the confidence interval satisfied

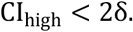

This criterion indicates that the observed variation across slices remains within a small multiple of the intrinsic variability, suggesting that the spatial distribution is effectively stable along that axis. Here, δ captures the baseline variability arising from within-slice noise, while the factor of 2 provides a tolerance margin that accounts for estimation uncertainty and variability across slices. In other words, allowing up to 2δ ensures that metabolites are not falsely classified as unstable due to minor fluctuations, while still requiring that between-slice variation remains comparable to the intrinsic noise level. Finally, to assess consistency across anatomical directions, results from the three axes were combined for each metabolite. Metabolites were categorized based on the number and combination of axes along which they were classified as stable, and overlap patterns were summarized using UpSet-style visualization. Overall, this analysis distinguishes true spatial variation from intrinsic noise and provides a principled basis for evaluating whether 2D sections can reliably approximate 3D spatial organization.

#### Two-group comparison using 2D data stratified by anatomical depth

When multiple 2D sections were available per group, statistical comparisons were performed at a depth-stratified level to account for the fact that sections acquired from different anatomical depths may reflect distinct spatial sampling of the tissue. Samples from each group were therefore partitioned into subsets corresponding to matched anatomical depth levels (depth 1, 2, and 3), and all analyses described below were applied independently within each depth stratum.

#### Depth-stratified global inter-region similarity comparison

For each sample within each depth stratum, an inter-region similarity matrix *S* ∈ *R*^R×R^ was computed using the optimal transport framework described above, where *R* denotes the number of anatomical regions. To obtain a scalar summary of global inter-region similarity for each sample, we extracted the upper triangular entries and vectorized them into a one-dimensional representation *v*. The sample-level summary statistic was then defined as the median of this vector, *s* = *media*n(*v*). This statistic captures the overall level of inter-region similarity while being robust to extreme values. Let 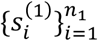 and 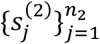 denote the sample-level summaries for two groups (e.g., WT and 5xFAD). The effect size was defined as the difference in group means:

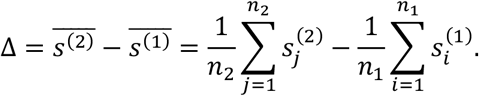

To assess statistical significance, we performed a permutation test under the null hypothesis of no group difference. Specifically, the pooled sample set 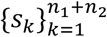 was randomly partitioned into two groups of sizes *n*_1_ and *n*_2_. For each permutation *b* = 1, …, *B*, the difference in group means was recomputed:

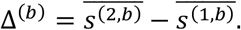

Repeating this procedure *B* = 1000 times yielded a null distribution {Δ^(b)^}. A two-sided permutation p-value was calculated as

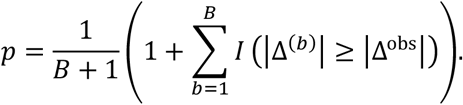

To quantify uncertainty in the estimated effect size, we performed bootstrap resampling within each group. For each bootstrap replicate *b*, samples were drawn with replacement from each group, and the effect size was recomputed as

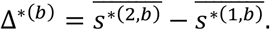

Repeating this procedure *B* = 10000 times produced a bootstrap distribution {Δ^∗(b)^}, from which a (1 − α) confidence interval was obtained using the percentile method:

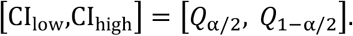

For visualization, results were presented using a Gardner–Altman plot, which simultaneously displays individual sample values, the permutation-based null distribution, and the bootstrap distribution of the effect size. This representation provides an integrated view of both statistical significance and effect size estimation.

#### Matrix-level pattern concordance analysis

To assess whether the overall pattern of inter-region metabolomic similarity was preserved within groups and altered between groups, we performed a matrix-level concordance analysis based on pairwise Spearman correlations of sample-specific similarity matrices. For each sample, the upper-triangular entries of the similarity matrix, excluding the diagonal, were then extracted and vectorized to represent the set of unique pairwise inter-region similarities. For any two samples, pattern concordance was quantified as the Spearman rank correlation between their vectorized similarity profiles, computed using only finite entries:

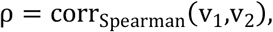

Within-group concordance was calculated separately for all pairwise comparisons among samples in group 1 and among samples in group 2. Between-group concordance was calculated for all pairwise comparisons between samples from different groups. The within-group concordance values from both groups were then pooled, and the observed test statistic was defined as the difference between the mean within-group and mean between-group concordance:

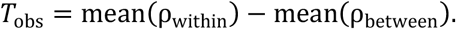

A larger value of *T*_obs_ indicates that similarity patterns are more consistent within groups than across groups. Statistical significance was assessed using a permutation test with *N* = 1000 permutations. Sample labels were randomly permuted while preserving group sizes, and the statistic *T* was recomputed for each permuted labeling to generate a null distribution 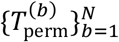. The one-sided permutation p-value was calculated as:

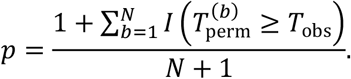

Results were visualized using boxplots and overlaid jittered points showing the distributions of within-group and between-group Spearman correlation coefficients.

#### Edge-wise similarity comparison using group-mean matrices

To compare condition-specific differences at the level of individual inter-region connections, we constructed group-mean similarity matrices for each condition by averaging the sample-specific similarity matrices within each group. The upper-triangular entries of the resulting group-mean matrices were then extracted and compared between conditions. Let *x* and *y* denote the vectorized upper-triangular entries from the two group-mean matrices. The concordance of edge-wise similarity values between groups was quantified using the Spearman rank correlation:

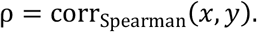

For visualization, these values were plotted in a scatter plot, where each point represents a single inter-region pair. This analysis provides a pairwise view of how inter-region similarity values are preserved or shifted between conditions.

#### Region-level connectivity change analysis

To characterize region-level patterns of metabolomic connectivity and evaluate the extent of condition-specific differences, we performed a region-level analysis based on sample-specific similarity matrices. For each sample, the mean connectivity of each anatomical region was defined as the average similarity between that region and all other regions, excluding self-similarity. Formally, for a given sample with similarity matrix S ∈ R^R × R^, the connectivity of region *r* was defined as

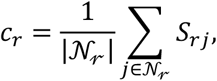

where 𝒩_𝓇_ = {*j* ≠ *r*}. This yielded a vector of region-level connectivity values for each sample, which were aggregated across samples to form group-level summaries. For each region, the difference in mean connectivity between groups was computed as

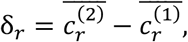

where 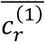 and 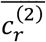 denote the average connectivity of region *r* across samples in group 1 and group 2, respectively. Statistical significance was assessed using a two-sided permutation test with *N* = 1000 permutations, in which sample labels were randomly reassigned while preserving group sizes to generate a null distribution of effect sizes. The permutation p-value for each region was computed as

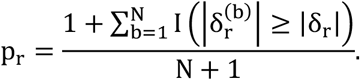

To account for multiple comparisons across regions, p-values were adjusted using the Benjamini–Hochberg false discovery rate (FDR) procedure. Regions were ranked according to the absolute magnitude of the observed effect size, and results were visualized using horizontal bar plots.

#### Metabolite Set Enrichment Analysis

Metabolite set enrichment analysis (MSEA) was performed to identify pathways showing coordinated metabolic differences between 5xFAD and WT coronal samples (n=3 for each group). For each sample, metabolite intensities were first averaged across all pixels belonging to the same sample, generating a sample-level metabolite abundance matrix. Pathway metabolite sets were constructed from the SMPDB pathway library, where each pathway was defined by its associated member metabolites. MSEA was then performed using a GSEA-style framework, in which metabolites were ranked according to the signal-to-noise statistic between the 5xFAD and WT groups. Pathways with fewer than 2 metabolites or more than 500 metabolites were excluded from the analysis. Statistical significance was assessed using 1,000 phenotype permutations, with a fixed random seed to ensure reproducibility. Enrichment results were summarized using p-values and normalized enrichment scores. Pathways with positive enrichment scores were interpreted as enriched in the 5xFAD group, whereas pathways with negative enrichment scores were interpreted as enriched in the WT group. The top enriched pathways were visualized using bar plots ranked by p-value.

#### General statistical methods

All animals and data points were included in statistical analyses. Data distribution was assumed to be normal but was not formally tested. Statistical analyses for behavioral and immunofluorescence data were performed using GraphPad Prism. Numerical data are presented as mean ± S.E.M. Comparisons were performed using one-way or two-way ANOVA or Student’s t-test as appropriate. P-values are indicated on each graph. Statistical parameters for each experiment are provided in the figures and figure legends.

### Individual metabolite analysis

For each metabolite, region-level abundance was computed by averaging TIC-normalized, log-transformed pixel intensities within each registered anatomical region of every section. Per-region between-group comparisons (5xFAD vs WT, HIF1α vs Control, or Neuronal AOX vs Control) were performed using a two-sided Wilcoxon rank-sum test on the pixel-level intensity distributions of matched regions, and the corresponding log2 fold change (log2FC) was computed from the region-mean intensities. For each comparison, p-values were adjusted across regions and across metabolites using the Benjamini–Hochberg false discovery rate (FDR) procedure. A metabolite was classified as predominantly up- or down-regulated across regions if more than two-thirds of the examined regions showed a consistent directional change at FDR < 0.05; metabolites that did not meet this criterion in either direction were classified as mixed (used in Figures 5O, 6M, and Supplementary Figures 16G, 19G, 23G, and 25G).

### Software and data availability

Image acquisition and primary spectral processing were performed in flexImaging v6.0 and SCiLS Lab v2024b (Bruker Daltonics). Three-dimensional MALDI volume reconstruction used the MetaVision3D pipeline, and multi-omics extraction used the Sami platform. Atlas registration used the QUINT workflow (QuickNII and VisuAlign) for manual alignment and a custom Python pipeline built on SimpleITK (v2.3) and tifffile for automatic registration. Coherence analysis was implemented in Python (v3.10) using AnnData, scanpy, NumPy, SciPy, scikit-learn, pandas, and the GeomLoss package for entropy-regularized optimal transport (Sinkhorn). Permutation tests, bootstrap analyses, and Spearman/Kolmogorov–Smirnov computations used SciPy. Behavioral video analysis used DeepLabCut and ANY-maze (Stoelting Co.). Whole-slide image quantification used HALO (Indica Labs). Proteomics data were processed with Proteome Discoverer v3.0 (Thermo Fisher Scientific) using the CHIMERYS search algorithm. Behavioral and immunofluorescence statistics were computed in GraphPad Prism. All raw MALDI imaging data, processed coherence matrices, and custom analysis code will be made publicly available at https://sunlabresources.rc.ufl.edu/ upon publication.

## References

1. Raichle, M.E., and Mintun, M.A. (2006). Brain work and brain imaging. Annu. Rev. Neurosci. 29, 449–476.

2. Vaishnavi, S.N., Vlassenko, A.G., Rundle, M.M., Snyder, A.Z., Mintun, M.A., and Raichle, M.E. (2010). Regional aerobic glycolysis in the human brain. Proc. Natl. Acad. Sci. USA 107, 17757–17762.

3. Vlassenko, A.G., Vaishnavi, S.N., Couture, L., Sacco, D., Shannon, B.J., Mach, R.H., Morris, J.C., Raichle, M.E., and Mintun, M.A. (2010). Spatial correlation between brain aerobic glycolysis and amyloid-beta deposition. Proc. Natl. Acad. Sci. USA 107, 17763–17767.

4. von Maydell, D., Wright, S.E., Pao, P., Staab, C., King, O., Spitaleri, A., Bonner, J.M., Liu, L., Yu, C.J., Chiu, C., et al. (2025). ABCA7 variants impact phosphatidylcholine and mitochondria in neurons. Nature 647, 462–471.

5. Mosconi, L. (2013). Glucose metabolism in normal aging and Alzheimer’s disease: methodological and physiological considerations for PET studies. Clin. Transl. Imaging 1, 217–233.

6. Mathys, H., Davila-Velderrain, J., Peng, Z., Gao, F., Mohammadi, S., Young, J.Z., Menon, M., He, L., Abdurrob, F., Jiang, X., et al. (2019). Single-cell transcriptomic analysis of Alzheimer’s disease. Nature 570, 332–337.

7. Mistur, R., Mosconi, L., Santi, S.D., Guzman, M., Li, Y., Tsui, W., and de Leon, M.J. (2009). Current Challenges for the Early Detection of Alzheimer’s Disease: Brain Imaging and CSF Studies. J. Clin. Neurol. 5, 153–166.

8. Minhas, P.S., Jones, J.R., Latif-Hernandez, A., Sugiura, Y., Durairaj, A.S., Wang, Q., Mhatre, S.D., Uenaka, T., Crapser, J., Conley, T., et al. (2024). Restoring hippocampal glucose metabolism rescues cognition across Alzheimer’s disease pathologies. Science 385, eabm6131.

9. Seeley, W.W., Crawford, R.K., Zhou, J., Miller, B.L., and Greicius, M.D. (2009). Neurodegenerative diseases target large-scale human brain networks. Neuron 62, 42–52.

10. Seguin, C., Sporns, O., and Zalesky, A. (2023). Brain network communication: concepts, models and applications. Nat. Rev. Neurosci. 24, 557–574.

11. Bassett, D.S., and Sporns, O. (2017). Network neuroscience. Nat. Neurosci. 20, 353–364.

12. Hawrylycz, M.J., Lein, E.S., Guillozet-Bongaarts, A.L., Shen, E.H., Ng, L., Miller, J.A., van de Lagemaat, L.N., Smith, K.A., Ebbert, A., Riley, Z.L., et al. (2012). An anatomically comprehensive atlas of the adult human brain transcriptome. Nature 489, 391–399.

13. Richiardi, J., Altmann, A., Milazzo, A., Chang, C., Chakravarty, M.M., Banaschewski, T., Barker, G.J., Bokde, A.L., Bromberg, U., Büchel, C., et al. (2015). Correlated gene expression supports synchronous activity in brain networks. Science 348, 1241–1244.

14. Yao, Z., van Velthoven, C.T.J., Kunst, M., Zhang, M., McMillen, D., Lee, C., Jung, W., Goldy, J., Abdelhak, A., Aitken, M., et al. (2023). A high-resolution transcriptomic and spatial atlas of cell types in the whole mouse brain. Nature 624, 317–332.

15. Zhang, M., Pan, X., Jung, W., Halpern, A.R., Eichhorn, S.W., Lei, Z., Cohen, L., Smith, K.A., Tasic, B., Yao, Z., et al. (2023). Molecularly defined and spatially resolved cell atlas of the whole mouse brain. Nature 624, 343–354.

16. Shi, H., He, Y., Zhou, Y., Huang, J., Maher, K., Wang, B., Tang, Z., Luo, S., Tan, P., Wu, M., et al. (2023). Spatial atlas of the mouse central nervous system at molecular resolution. Nature 622, 552–561.

17. Ma, M., Yu, Q., Delafield, D.G., Cui, Y., Li, Z., Li, M., Wu, W., Shi, X., Westmark, P.R., Gutierrez, A., et al. (2024). On-Tissue Spatial Proteomics Integrating MALDI-MS Imaging with Shotgun Proteomics Reveals Soy Consumption-Induced Protein Changes in a Fragile X Syndrome Mouse Model. ACS Chem. Neurosci. 15, 119–133.

18. Sharma, K., Hansen, J., Susztak, K., Eberlin, L., Anderton, C.R., Alexandrov, T., and Iyengar, R. (2025). Spatial metabolomics and multiomics integration for breakthroughs in precision medicine for kidney disease. Nat. Rev. Nephrol. 22, 152–164.

19. Nwabufo, C.K., and Aigbogun, O.P. (2022). Potential application of mass spectrometry imaging in pharmacokinetic studies. Xenobiotica 52, 811–827.

20. Lee, Y.-R., Kaya, I., Wik, E., Baijnath, S., Lodén, H., Nilsson, A., Zhang, X., Sehlin, D., Syvänen, S., Svenningsson, P., and Andrén, P.E. (2025). Comprehensive Approach for Sequential MALDI-MSI Analysis of Lipids, N-Glycans, and Peptides in Fresh-Frozen Rodent Brain Tissues. Anal. Chem. 97, 1338–1346.

21. Kaya, I., Schembri, L.S., Nilsson, A., Shariatgorji, R., Baijnath, S., Zhang, X., Bezard, E., Svenningsson, P., Odell, L.R., and Andrén, P.E. (2023). On-Tissue Chemical Derivatization for Comprehensive Mapping of Brain Carboxyl and Aldehyde Metabolites by MALDI-MS Imaging. J. Am. Soc. Mass Spectrom. 34, 836–846.

22. Djambazova, K.V., Dufresne, M., Migas, L.G., Kruse, A.R.S., Van de Plas, R., Caprioli, R.M., and Spraggins, J.M. (2023). MALDI TIMS IMS of Disialoganglioside Isomers─GD1a and GD1b in Murine Brain Tissue. Anal. Chem. 95, 1176–1183.

23. Kaya, I., Nilsson, A., Luptaková, D., He, Y., Vallianatou, T., Bjärterot, P., Svenningsson, P., Bezard, E., and Andrén, P.E. (2023). Spatial lipidomics reveals brain region-specific changes of sulfatides in an experimental MPTP Parkinson’s disease primate model. npj Parkinsons Dis. 9, 118.

24. Clarke, H.A., Ma, X., Shedlock, C.J., Medina, T., Hawkinson, T.R., Wu, L., Ribas, R.A., Keohane, S., Ravi, S., Bizon, J.L., et al. (2025). Spatial mapping of the brain metabolome lipidome and glycome. Nat. Commun. 16, 4373.

25. Pádua, M.S., Guil-Guerrero, J.L., and Lopes, P.A. (2024). Behaviour Hallmarks in Alzheimer’s Disease 5xFAD Mouse Model. Int. J. Mol. Sci. 25, 6766.

26. Kosel, F., Pelley, J.M., and Franklin, T.B. (2020). Behavioural and psychological symptoms of dementia in mouse models of Alzheimer’s disease-related pathology. Neurosci. Biobehav. Rev. 112, 634–647.

27. Ma, X., Shedlock, C.J., Medina, T., Ribas, R.A., Clarke, H.A., Hawkinson, T.R., Dande, P.K., Golamari, H.K.R., Wu, L., Ziani, B.E., et al. (2025). AI-driven framework to map the brain metabolome in three dimensions. Nat. Metab. 7, 842–853.

28. Bunne, C., Schiebinger, G., Krause, A., Regev, A., and Cuturi, M. (2024). Optimal transport for single-cell and spatial omics. Nat. Rev. Methods Primers 4, 58.

29. Schiebinger, G., Shu, J., Tabaka, M., Cleary, B., Subramanian, V., Solomon, A., Gould, J., Liu, S., Lin, S., Berube, P., et al. (2019). Optimal-Transport Analysis of Single-Cell Gene Expression Identifies Developmental Trajectories in Reprogramming. Cell 176, 928-943.e22.

30. Semenza, G.L. (2013). HIF-1 mediates metabolic responses to intratumoral hypoxia and oncogenic mutations. J. Clin. Invest. 123, 3664–3671.

31. Martínez-Reyes, I., Cardona, L.R., Kong, H., Vasan, K., McElroy, G.S., Werner, M., Kihshen, H., Reczek, C.R., Weinberg, S.E., Gao, P., et al. (2020). Mitochondrial ubiquinol oxidation is necessary for tumour growth. Nature 585, 288–292.

32. Humphrey, D.M., Parsons, R.B., Ludlow, Z.N., Riemensperger, T., Esposito, G., Verstreken, P., Jacobs, H.T., Birman, S., and Hirth, F. (2012). Alternative oxidase rescues mitochondria-mediated dopaminergic cell loss in Drosophila. Hum. Mol. Genet. 21, 2698–2712.

33. Kemppainen, K.K., Rinne, J., Sriram, A., Lakanmaa, M., Zeb, A., Tuomela, T., Popplestone, A., Singh, S., Sanz, A., Rustin, P., and Jacobs, H.T. (2014). Expression of alternative oxidase in Drosophila ameliorates diverse phenotypes due to cytochrome oxidase deficiency. Hum. Mol. Genet. 23, 2078–2093.

34. Szibor, M., Schenkl, C., Barsottini, M.R.O., Young, L., and Moore, A.L. (2022). Targeting the alternative oxidase (AOX) for human health and food security, a pharmaceutical and agrochemical target or a rescue mechanism? Biochem. J. 479, 1337–1359.

35. Diaz, F., Garcia, S., Padgett, K.R., and Moraes, C.T. (2012). A defect in the mitochondrial complex III, but not complex IV, triggers early ROS-dependent damage in defined brain regions. Hum. Mol. Genet. 21, 5066–5077.

36. Barnett, D., Zimmer, T.S., Booraem, C., Palaguachi, F., Meadows, S.M., Xiao, H., Wong, M.Y., Luo, W., Gan, L., Chouchani, E.T., et al. (2025). Mitochondrial complex III-derived ROS amplify immunometabolic changes in astrocytes and promote dementia pathology. Nat. Metab. 7, 2300–2323.

37. Goyal, M.S., Blazey, T., Metcalf, N.V., McAvoy, M.P., Strain, J.F., Rahmani, M., Durbin, T.J., Xiong, C., Benzinger, T.L., Morris, J.C., et al. (2023). Brain aerobic glycolysis and resilience in Alzheimer disease. Proc. Natl. Acad. Sci. USA 120, e2212256120.

38. Lucchetti, F., Céléreau, E., Steullet, P., Alemán-Gómez, Y., Hagmann, P., Klauser, A., and Klauser, P. (2025). Constructing the human brain metabolic connectome with MR spectroscopic imaging reveals cerebral biochemical organization. Nat. Commun. 16, 11344.

39. Pellerin, L., and Magistretti, P.J. (2012). Sweet sixteen for ANLS. J. Cereb. Blood Flow Metab. 32, 1152–1166.

40. Bullmore, E., and Sporns, O. (2012). The economy of brain network organization. Nat. Rev. Neurosci. 13, 336–349.

41. Buckner, R.L., Sepulcre, J., Talukdar, T., Krienen, F.M., Liu, H., Hedden, T., Andrews-Hanna, J.R., Sperling, R.A., and Johnson, K.A. (2009). Cortical hubs revealed by intrinsic functional connectivity: mapping, assessment of stability, and relation to Alzheimer’s disease. J. Neurosci. 29, 1860–1873.

42. Pascoal, T.A., Mathotaarachchi, S., Kang, M.S., Mohaddes, S., Shin, M., Park, A.Y., Parent, M.J., Benedet, A.L., Chamoun, M., Therriault, J., et al. (2019). Aβ-induced vulnerability propagates via the brain’s default mode network. Nat. Commun. 10, 2353.

43. Therriault, J., Pascoal, T.A., Savard, M., Mathotaarachchi, S., Benedet, A.L., Chamoun, M., Tissot, C., Lussier, F.Z., Rahmouni, N., Stevenson, J., et al. (2022). Intrinsic connectivity of the human brain provides scaffold for tau aggregation in clinical variants of Alzheimer’s disease. Sci. Transl. Med. 14, eabc8693.

44. Cope, T.E., Rittman, T., Borchert, R.J., Jones, P.S., Vatansever, D., Allinson, K., Passamonti, L., Vazquez Rodriguez, P., Bevan-Jones, W.R., O’Brien, J.T., and Rowe, J.B. (2018). Tau burden and the functional connectome in Alzheimer’s disease and progressive supranuclear palsy. Brain 141, 550–567.

45. Wang, G., van den Berg, B.M., Kostidis, S., Pinkham, K., Jacobs, M.E., Liesz, A., Giera, M., and Rabelink, T.J. (2025). Spatial quantitative metabolomics enables identification of remote and sustained ipsilateral cortical metabolic reprogramming after stroke. Nat. Metab. 7, 1791–1800.

